## Additional File 1 for "Synthetic evolution of herbicide resistance using a T7 RNAP-based random DNA base editor"

**Additional File 1:**

**Figure S1:** Amplicon deep sequencing analysis of the *GFP* target in *Nicotiana benthamiana*.

**Table S1:** List of sequences used in this study

**Sequences S1:** DNA sequence of the vector attL1_3xFLAG_AID_T7_UGI _attL2_pUC57

**Sequences S2:** DNA sequence of the vector pEditor: **p35S-T7pol** (p35S_3xFLAG_T7_UGI _pK2GW7) (two-vector system)

**Sequences S3:** DNA sequence of the vector pEditor: **p35S-AID-T7pol** (p35S_3xFLAG_AID_T7_UGI _pK2GW7) (two-vector system)

**Sequences S4:** DNA sequence of the vector pTarget: **p35S-pT7-GFP** (p35S_pT7_GFP_NOSterm) (two-vector system)

**Sequences S5:** DNA sequence of the vector pTarget: **p35S-pT7-ALS** (p35S_pT7_ALS_T7term_polA) (two-vector system)

**Sequences S6:** DNA sequence of the single-vector system **p35S-pT7-ALS/pUBI-AID-T7pol** (p35S_pT7_ALS_T7term_polA_pUBI_AID_T7pol_UGI_NOSterm)

**p35s_GFP_F7R1_Data_1**

|  | **Wild Type** | **Insertions** | **Deletions** | **Indel Rate (%)** | **Substitution Rate (%)** |
| --- | --- | --- | --- | --- | --- |
| Experimental samples | 216,471 | 41 | 219 | 0.12 | 6.76 (14,659) |
| Mock-treated samples | 232,560 | 35 | 201 | 0.10 | N.A. |

|  | **Wild Type** | **Insertions** | **Deletions** | **Indel Rate (%)** | **Substitution Rate (%)** |
| --- | --- | --- | --- | --- | --- |
| Experimental samples | 218,265 | 26 | 181 | 0.09 | 6.37 (13,914) |
| Mock-treated samples | 226,511 | 31 | 180 | 0.09 | N.A. |

**p35s_GFP_F7R1_Data_2**

**p35s_GFP_FR2_Data_1**

|  | **Wild Type** | **Insertions** | **Deletions** | **Indel Rate (%)** | **Substitution Rate (%)** |
| --- | --- | --- | --- | --- | --- |
| Experimental samples | 225,572 | 83 | 286 | 0.16 | 9.13 (20,628) |
| Mock-treated samples | 229,871 | 71 | 255 | 0.14 | N.A. |

**p35s_GFP_FR2_Data_2**

|  | **Wild Type** | **Insertions** | **Deletions** | **Indel Rate (%)** | **Substitution Rate (%)** |
| --- | --- | --- | --- | --- | --- |
| Experimental samples | 231,886 | 55 | 299 | 0.15 | 8.47 (19,664) |
| Mock-treated samples | 230,399 | 70 | 258 | 0.14 | N.A. |

**Figure S1:** Amplicon deep sequencing analysis of the *GFP* target in *Nicotiana benthamiana*. The data were analyzed using the web tool CRISPR-sub (Hwang et al., 2020), and mock-treated reads were subtracted from experimental sample reads to calculate the editing efficiency. The table shows the number of reads.

**Table S1: List of sequences used in this study**

| **Name of the primer** | **Sequence (5' - 3')** | **Purpose** |
| --- | --- | --- |
| SphI_AID_T7_F | CCGCATGCCCCGGGatggacagcctcttgatgaaccg | Cloning of pEditor |
| NSiI_XbaI_AID_T7_R | CCTCTAGAATGCATgactttcctcttcttcttgggagaaccaccagagag | Cloning of pEditor |
| SbfI_XbaI_T7_pro_F | ggtaatacgactcactataggGAGACCACAACGGTTTCCCTAGAAATAATTTTGTTTAACTTTAAGaaggagATATAt | Cloning of T7 promoter upstream of *GFP* |
| SbfI_XbaI_T7_pro_R | ctagaTATATctccttCTTAAAGTTAAACAAAATTATTTCTAGGGAAACCGTTGTGGTCTCcctatagtgagtcgtattacctgca | Cloning of T7 promoter upstream of *GFP* |
| SacI_T7_Ter_F | cactagcataaccccttggggcctctaaacgggtcttgaggggttttttggagct | Cloning of T7 terminator downstream of *GFP* |
| SacI_T7_Ter_R | ccaaaaaacccctcaagacccgtttagaggccccaaggggttatgctagtgagct | Cloning of T7 terminator downstream of *GFP* |
| 35sP_HinD_F | ccaagctttgagacttttcaacaaagggtaatatcggg | Cloning of 35S promoter |
| 35sP_PstI_HinD_R | ttaagcttctgcagtcagcgtgtcctctccaaatg | Cloning of 35S promoter |
| T7_PT_F_1 | agctttaatacgactcactataggGAGACCACAACGGTTTCCCTAGAAATAATTTTGTTTAACTTTAAGaaggagATATAGGGCCC | T7-promoter_RBS_MCS_T7-terminator_CaMV poly(A)_fragment_1 |
| T7_PT_R_1 | CCTAGGGGGCCCTATATctccttCTTAAAGTTAAACAAAATTATTTCTAGGGAAACCGTTGTGGTCTCcctatagtgagtcgtattaa | T7-promoter_RBS_MCS_T7-terminator_CaMV poly(A)_fragment_1 |
| T7_PT_F_2 | CCTAGGGGTACCTCTAGAactagcataaccccttggggcctctaaacgggtcttgaggggttttttgtttctccataataatgtgt | T7-promoter_RBS_MCS_T7-terminator_CaMV poly(A)_fragment_2 |
| T7_PT_R_2 | ctactcacacattattatggagaaacaaaaaacccctcaagacccgtttagaggccccaaggggttatgctagtTCTAGAGGTACC | T7-promoter_RBS_MCS_T7-terminator_CaMV poly(A)_fragment_2 |
| T7_PT_F_3 | gagtagttcccagataagggaattagggttcctatagggtttcgctcatgtgttgagcatataagaaacccttagtatgtatttgt | T7-promoter_RBS_MCS_T7-terminator_CaMV poly(A)_fragment_3 |
| T7_PT_R_3 | acaaatacaaatacatactaagggtttcttatatgctcaacacatgagcgaaaccctataggaaccctaattcccttatctgggaa | T7-promoter_RBS_MCS_T7-terminator_CaMV poly(A)_fragment_3 |
| T7_PT_F_4 | atttgtaaaatacttctatcaataaaatttctaattcctaaaaccaaaatccagtactaaaatccagatctgcatgcctgca | T7-promoter_RBS_MCS_T7-terminator_CaMV poly(A)_fragment_4 |
| T7_PT_R_4 | ggcatgcagatctggattttagtactggattttggttttaggaattagaaattttattgatagaagtatttt | T7-promoter_RBS_MCS_T7-terminator_CaMV poly(A)_fragment_4 |
| HindIII_T7_PT_F | CCaagctttaatacgactcactataggG | Cloning of T7-promoter_RBS_MCS_T7-terminator_CaMV poly(A) in pRGEB32 |
| Sbf_T7_PT_R | TTcctgcaggcatgcagatc | Cloning of T7-promoter_RBS_MCS_T7-terminator_CaMV poly(A) in pRGEB32 |
| ApaI_AvrII_ALS_F | AAGGGCCCCCTAGGATGGCTACGACCGCCGCGGC | Cloning of *ALS* in pRGEB32 driven by p35S |
| XbaI_KpnI_ALS_R | GGTCTAGAGGTACCTTAATACACAGTCCTGCCATCACCATCCAGG | Cloning of *ALS* in pRGEB32 driven by p35S |
| GFP_F7 | gccgaattcagtaaaggagaag | Amplicon deep sequencing of *GFP* target |
| GFP_R1 | ctccttgaaatcgattcccttaag | Amplicon deep sequencing of *GFP* target |
| PGD_GFP_F | caacgtctatatcatggccgac | Amplicon deep sequencing of *GFP* target |
| GFP_R2 | ctcatcatgtttgtatagttcatccatg | Amplicon deep sequencing of *GFP* target |

**Sequences S1:** DNA sequence of the vector attL1_3xFLAG_AID_T7_UGI _attL2_pUC57

CCAATGATCAAATAATGATTTTATTTTGACTGATAGTGACCTGTTCGTTGCAACAAATTGATGAGCAATGCTTTTTTATAATGCCAACTTTGTACAAAAAAGCAGGCTTCGAAGGAGATAGAACCAATTCTCTAAGGAAATACTTAACCATGGACTATAAGGACCACGACGGAGACTACAAGGATCATGATATTGATTACAAAGACGATGACGATAAGATGGCCCCAAAGAAGAAGCGGAAGGTGGGTATCCACGGAGTCCCAGCAGCCGCATGCCCCGGGATGGACAGCCTCTTGATGAACCGGAGGGAGTTTCTTTACCAATTCAAAAATGTCCGCTGGGCTAAGGGTCGGCGTGAGACCTACCTGTGCTACGTAGTGAAGAGGCGTGACAGTGCTACATCCTTTTCACTGGACTTTGGTTATCTTCGCAATAAGAACGGCTGCCACGTGGAATTGCTCTTCCTCCGCTACATCTCGGACTGGGACCTAGACCCTGGCCGCTGCTACCGCGTCACCTGGTTCATCTCCTGGAGCCCCTGCTACGACTGTGCCCGACATGTGGCCGACTTTCTGCGAGGGAACCCCAACCTCAGTCTGAGGATCTTCACCGCGCGCCTCTACTTCTGTGAGGACCGCAAGGCTGAGCCCGAGGGGCTGCGGCGGCTGCACCGCGCCGGGGTGCAAATAGCCATCATGACCTTCAAAGATTATTTTTACTGCTGGAATACTTTTGTAGAAAACCACGGAAGAACTTTCAAAGCCTGGGAAGGGCTGCATGAAAATTCAGTTCGTCTCTCCAGACAGCTTCGGCGCATCCTTTTGCCCCTGTATGAGGTTGATGACTTACGAGACGCATTTCGTACTAGCGGCAGCGAGACTCCCGGGACCTCAGAGTCCGCCACACCCGAAAGTAACACCATCAACATTGCTAAGAACGACTTCTCAGACATAGAGCTCGCGGCTATTCCGTTCAACACCCTGGCTGACCACTACGGCGAGAGACTCGCTAGGGAGCAGCTGGCGTTGGAGCATGAATCCTACGAGATGGGCGAGGCTAGGTTCCGCAAGATGTTCGAGCGACAATTGAAGGCAGGGGAGGTGGCGGACAACGCTGCCGCCAAGCCCCTGATCACAACCTTGCTGCCCAAAATGATCGCGCGGATCAACGATTGGTTTGAGGAGGTTAAGGCAAAACGGGGCAAACGCCCGACCGCATTTCAATTCCTCCAAGAAATCAAGCCTGAGGCTGTTGCCTACATCACTATCAAGACGACACTGGCGTGTCTCACAAGCGCCGACAACACCACCGTGCAAGCCGTCGCCAGCGCCATCGGGCGGGCAATTGAGGATGAGGCACGGTTTGGTAGGATCCGAGACCTGGAAGCGAAGCACTTCAAGAAGAACGTGGAAGAGCAGTTGAACAAACGCGTCGGCCACGTGTATAAAAAGGCTTTCATGCAGGTGGTGGAGGCCGATATGCTCAGTAAGGGGCTGCTTGGGGGGGAGGCGTGGTCATCCTGGCACAAGGAGGATAGCATTCACGTGGGGGTCCGATGTATCGAGATGCTGATAGAGAGCACCGGAATGGTCTCCCTCCATCGCCAGAACGCTGGGGTCGTAGGGCAGGACTCCGAGACTATTGAGCTGGCCCCCGAGTATGCCGAAGCAATCGCTACACGCGCAGGTGCACTGGCTGGGATAAGCCCTATGTTTCAGCCCTGCGTAGTGCCTCCAAAGCCATGGACCGGCATCACAGGGGGTGGCTATTGGGCCAACGGTAGGCGGCCTCTGGCCCTGGTACGCACGCACAGCAAGAAGGCGCTCATGCGCTATGAAGACGTTTACATGCCCGAGGTTTACAAGGCGATCAATATCGCGCAGAACACCGCCTGGAAAATCAATAAGAAGGTGTTGGCGGTCGCAAACGTGATTACCAAGTGGAAGCATTGCCCAGTCGAGGACATACCCGCCATAGAACGCGAAGAGCTGCCGATGAAGCCGGAAGACATTGATATGAACCCCGAGGCCCTCACCGCGTGGAAAAGAGCCGCAGCCGCCGTATACAGGAAGGATAAAGCGCGCAAGTCCCTACGCATAAGCCTCGAGTTTATGCTGGAACAGGCCAACAAGTTCGCCAACCACAAAGCTATCTGGTTCCCCTACAACATGGACTGGAGAGGGAGGGTCTACGCCGTCAGCATGTTCAATCCCCAGGGCAACGACATGACGAAGGGCCTTCTGACATTGGCAAAGGGGAAGCCTATCGGAAAGGAGGGGTACTACTGGCTCAAGATCCACGGCGCCAACTGCGCGGGAGTGGACAAGGTTCCATTTCCCGAGCGAATTAAGTTCATCGAGGAAAACCACGAAAACATTATGGCGTGCGCTAAATCCCCCCTCGAGAACACATGGTGGGCCGAGCAAGACTCCCCGTTCTGTTTTTTGGCATTCTGCTTTGAGTACGCCGGTGTGCAGCACCATGGCCTCTCATACAACTGTTCCCTGCCCCTGGCCTTCGACGGAAGTTGCAGTGGGATTCAACATTTCAGCGCAATGTTGCGGGACGAGGTCGGTGGCAGGGCCGTTAACCTGCTCCCTTCCGAAACGGTGCAGGACATCTACGGAATCGTGGCAAAAAAGGTAAACGAGATCCTGCAAGCGGATGCCATCAACGGGACGGACAATGAGGTCGTTACGGTGACAGACGAAAATACTGGGGAAATAAGCGAAAAGGTCAAGCTGGGGACCAAAGCACTCGCGGGTCAGTGGCTCGCCTACGGGGTGACACGCTCCGTCACCAAGAGAAGCGTGATGACCCTCGCGTACGGTTCAAAAGAATTCGCCTTCCGCCAGCAAGTGCTGGAGGACACCATCCAGCCGGCGATTGACTCCGGGAAGGGTCTCATGTTTACCCAGCCGAACCAGGCCGCAGGGTACATGGCCAAACTGATCTGGGAAAGCGTTAGCGTCACAGTGGTCGCCGCGGTTGAGGCGATGAATTGGCTGAAGAGCGCGGCAAAGCTCCTCGCCGCTGAGGTGAAGGACAAAAAGACCGGCGAAATCCTGCGCAAGCGCTGCGCCGTCCACTGGGTCACGCCGGATGGATTCCCCGTCTGGCAGGAGTACAAGAAGCCCATCCGAACCCGGCTCAACTTGATGTTCCTTGGCCAGTTTCGCCTGCAGCCCACGATAAACACCAACAAAGACAGCGAGATCGACGCCCACAAGCAGGAGAGCGGCATCGCGCCCAACTTCGTGCACAGTCAGGACGGGTCCCATCTGCGGAAAACTGTTGTGTGGGCTCACGAGAAGTACGGCATTGAGAGCTTCGCCCTGATACACGACAGCTTCGGGACCATACCAGCGGACGCAGCGAACCTGTTCAAAGCCGTGCGGGAAACAATGGTCGACACCTACGAAAGCTGCGACGTACTGGCAGACTTCTATGACCAATTCGCCGACCAGCTTCACGAGTCACAGCTCGACAAGATGCCCGCTCTGCCCGCGAAAGGCAACCTGAATTTGCGCGACATCCTTGAGAGCGATTTTGCGTTCGCCTCTGGTGGTTCTACTAATCTGTCAGATATTATTGAAAAGGAGACCGGTAAGCAACTGGTTATCCAGGAATCCATCCTCATGCTCCCAGAGGAGGTGGAAGAAGTCATTGGGAACAAGCCGGAAAGCGATATACTCGTGCACACCGCCTACGACGAGAGCACCGACGAGAATGTCATGCTTCTGACTAGCGACGCCCCTGAATACAAGCCTTGGGCTCTGGTCATACAGGATAGCAACGGTGAGAACAAGATTAAGATGCTCTCTGGTGGTTCTCCCAAGAAGAAGAGGAAAGTCATGCATTAGACCCAGCTTTCTTGTACAAAGTTGGCATTATAAGAAAGCATTGCTTATCAATTTGTTGCAACGAACAGGTCACTATCAGTCAAAATAAAATCATTATTTGATCGGAAAGAACATGTGAGCAAAAGGCCAGCAAAAGGCCAGGAACCGTAAAAAGGCCGCGTTGCTGGCGTTTTTCCATAGGCTCCGCCCCCCTGACGAGCATCACAAAAATCGACGCTCAAGTCAGAGGTGGCGAAACCCGACAGGACTATAAAGATACCAGGCGTTTCCCCCTGGAAGCTCCCTCGTGCGCTCTCCTGTTCCGACCCTGCCGCTTACCGGATACCTGTCCGCCTTTCTCCCTTCGGGAAGCGTGGCGCTTTCTCATAGCTCACGCTGTAGGTATCTCAGTTCGGTGTAGGTCGTTCGCTCCAAGCTGGGCTGTGTGCACGAACCCCCCGTTCAGCCCGACCGCTGCGCCTTATCCGGTAACTATCGTCTTGAGTCCAACCCGGTAAGACACGACTTATCGCCACTGGCAGCAGCCACTGGTAACAGGATTAGCAGAGCGAGGTATGTAGGCGGTGCTACAGAGTTCTTGAAGTGGTGGCCTAACTACGGCTACACTAGAAGAACAGTATTTGGTATCTGCGCTCTGCTGAAGCCAGTTACCTTCGGAAAAAGAGTTGGTAGCTCTTGATCCGGCAAACAAACCACCGCTGGTAGCGGTGGTTTTTTTGTTTGCAAGCAGCAGATTACGCGCAGAAAAAAAGGATCTCAAGAAGATCCTTTGATCTTTTCTACGGGGTCTGACGCTCAGTGGAACGAAAACTCACGTTAAGGGATTTTGGTCATGAGATTATCAAAAAGGATCTTCACCTAGATCCTTTTAAATTAAAAATGAAGTTTTAAATCAATCTAAAGTATATATGAGTAAACTTGGTCTGACAGTTACCAATGCTTAATCAGTGAGGCACCTATCTCAGCGATCTGTCTATTTCGTTCATCCATAGTTGCCTGACTCCCCGTCGTGTAGATAACTACGATACGGGAGGGCTTACCATCTGGCCCCAGTGCTGCAATGATACCGCGAGACCCACGCTCACCGGCTCCAGATTTATCAGCAATAAACCAGCCAGCCGGAAGGGCCGAGCGCAGAAGTGGTCCTGCAACTTTATCCGCCTCCATCCAGTCTATTAATTGTTGCCGGGAAGCTAGAGTAAGTAGTTCGCCAGTTAATAGTTTGCGCAACGTTGTTGCCATTGCTACAGGCATCGTGGTGTCACGCTCGTCGTTTGGTATGGCTTCATTCAGCTCCGGTTCCCAACGATCAAGGCGAGTTACATGATCCCCCATGTTGTGCAAAAAAGCGGTTAGCTCCTTCGGTCCTCCGATCGTTGTCAGAAGTAAGTTGGCCGCAGTGTTATCACTCATGGTTATGGCAGCACTGCATAATTCTCTTACTGTCATGCCATCCGTAAGATGCTTTTCTGTGACTGGTGAGTACTCAACCAAGTCATTCTGAGAATAGTGTATGCGGCGACCGAGTTGCTCTTGCCCGGCGTCAATACGGGATAATACCGCGCCACATAGCAGAACTTTAAAAGTGCTCATCATTGGAAAACGTTCTTCGGGGCGAAAACTCTCAAGGATCTTACCGCTGTTGAGATCCAGTTCGATGTAACCCACTCGTGCACCCAACTGATCTTCAGCATCTTTTACTTTCACCAGCGTTTCTGGGTGAGCAAAAACAGGAAGGCAAAATGCCGCAAAAAAGGGAATAAGGGCGACACGGAAATGTTGAATACTCATACTCTTCCTTTTTCAATATTATTGAAGCATTTATCAGGGTTATTGTCTCATGAGCGGATACATATTTGAATGTATTTAGAAAAATAAACAAATAGGGGTTCCGCGCACATTTCCCCGAAAAGTGCCACCTGACGTC

**Sequences S2:** DNA sequence of the vector pEditor: **p35S-T7pol** (p35S_3xFLAG_T7_UGI _pK2GW7) (two-vector system)

CCGGCATATGGTCGACTAGAGCCAAGCTGATCTCCTTTGCCCCGGAGATCACCATGGACGACTTTCTCTATCTCTACGATCTAGGAAGAAAGTTCGACGGAGAAGGTGACGATACCATGTTCACCACCGATAATGAGAAGATTAGCCTCTTCAATTTCAGAAAGAATGCTGACCCACAGATGGTTAGAGAGGCCTACGCGGCAGGTCTCATCAAGACGATCTACCCGAGTAATAATCTCCAGGAGATCAAATACCTTCCCAAGAAGGTTAAAGATGCAGTCAAAAGATTCAGGACTAACTGCATCAAGAACACAGAGAAAGATATATTTCTCAAGATCAGAAGTACTATTCCAGTATGGACGATTCAAGGCTTGCTTCATAAACCAAGGCAAGTAATAGAGATTGGAGTCTCTAAGAAAGTAGTTCCTACTGAATCAAAGGCCATGGAGTCAAAAATTCAGATCGAGGATCTAACAGAACTCGCCGTGAAGACTGGCGAACAGTTCATACAGAGTCTTTTACGACTCAATGACAAGAAGAAAATCTTCGTCAACATGGTGGAGCACGACACTCTCGTCTACTCCAAGAATATCAAAGATACAGTCTCAGAAGACCAAAGGGCTATTGAGACTTTTCAACAAAGGGTAATATCGGGAAACCTCCTCGGATTCCATTGCCCAGCTATCTGTCACTTCATCAAAAGGACAGTAGAAAAGGAAGGTGGCACCTACAAATGCCATCATTGCGATAAAGGAAAGGCTATCGTTCAAGATGCCTCTGCCGACAGTGGTCCCAAAGATGGACCCCCACCCACGAGGAGCATCGTGGAAAAAGAAGACGTTCCAACCACGTCTTCAAAGCAAGTGGATTGATGTGATATCTCCACTGACGTAAGGGATGACGCACAATCCCACTATCCTTCGCAAGACCCTTCCTCTATATAAGGAAGTTCATTTCATTTGGAGAGGACTCCGGTATTTTTACAACAATACCACAACAAAACAAACAACAAACAACATTACAATTTACTATTCTAGTCGACCTGCAGGCGGCCGCACTAGTGATATCACAAGTTTGTACAAAAAAGCAGGCTTCGAAGGAGATAGAACCAATTCTCTAAGGAAATACTTAACCATGGACTATAAGGACCACGACGGAGACTACAAGGATCATGATATTGATTACAAAGACGATGACGATAAGATGGCCCCAAAGAAGAAGCGGAAGGTGGGTATCCACGGAGTCCCAGCAGCCGCATGCCCCGGGACCTCAGAGTCCGCCACACCCGAAAGTAACACCATCAACATTGCTAAGAACGACTTCTCAGACATAGAGCTCGCGGCTATTCCGTTCAACACCCTGGCTGACCACTACGGCGAGAGACTCGCTAGGGAGCAGCTGGCGTTGGAGCATGAATCCTACGAGATGGGCGAGGCTAGGTTCCGCAAGATGTTCGAGCGACAATTGAAGGCAGGGGAGGTGGCGGACAACGCTGCCGCCAAGCCCCTGATCACAACCTTGCTGCCCAAAATGATCGCGCGGATCAACGATTGGTTTGAGGAGGTTAAGGCAAAACGGGGCAAACGCCCGACCGCATTTCAATTCCTCCAAGAAATCAAGCCTGAGGCTGTTGCCTACATCACTATCAAGACGACACTGGCGTGTCTCACAAGCGCCGACAACACCACCGTGCAAGCCGTCGCCAGCGCCATCGGGCGGGCAATTGAGGATGAGGCACGGTTTGGTAGGATCCGAGACCTGGAAGCGAAGCACTTCAAGAAGAACGTGGAAGAGCAGTTGAACAAACGCGTCGGCCACGTGTATAAAAAGGCTTTCATGCAGGTGGTGGAGGCCGATATGCTCAGTAAGGGGCTGCTTGGGGGGGAGGCGTGGTCATCCTGGCACAAGGAGGATAGCATTCACGTGGGGGTCCGATGTATCGAGATGCTGATAGAGAGCACCGGAATGGTCTCCCTCCATCGCCAGAACGCTGGGGTCGTAGGGCAGGACTCCGAGACTATTGAGCTGGCCCCCGAGTATGCCGAAGCAATCGCTACACGCGCAGGTGCACTGGCTGGGATAAGCCCTATGTTTCAGCCCTGCGTAGTGCCTCCAAAGCCATGGACCGGCATCACAGGGGGTGGCTATTGGGCCAACGGTAGGCGGCCTCTGGCCCTGGTACGCACGCACAGCAAGAAGGCGCTCATGCGCTATGAAGACGTTTACATGCCCGAGGTTTACAAGGCGATCAATATCGCGCAGAACACCGCCTGGAAAATCAATAAGAAGGTGTTGGCGGTCGCAAACGTGATTACCAAGTGGAAGCATTGCCCAGTCGAGGACATACCCGCCATAGAACGCGAAGAGCTGCCGATGAAGCCGGAAGACATTGATATGAACCCCGAGGCCCTCACCGCGTGGAAAAGAGCCGCAGCCGCCGTATACAGGAAGGATAAAGCGCGCAAGTCCCTACGCATAAGCCTCGAGTTTATGCTGGAACAGGCCAACAAGTTCGCCAACCACAAAGCTATCTGGTTCCCCTACAACATGGACTGGAGAGGGAGGGTCTACGCCGTCAGCATGTTCAATCCCCAGGGCAACGACATGACGAAGGGCCTTCTGACATTGGCAAAGGGGAAGCCTATCGGAAAGGAGGGGTACTACTGGCTCAAGATCCACGGCGCCAACTGCGCGGGAGTGGACAAGGTTCCATTTCCCGAGCGAATTAAGTTCATCGAGGAAAACCACGAAAACATTATGGCGTGCGCTAAATCCCCCCTCGAGAACACATGGTGGGCCGAGCAAGACTCCCCGTTCTGTTTTTTGGCATTCTGCTTTGAGTACGCCGGTGTGCAGCACCATGGCCTCTCATACAACTGTTCCCTGCCCCTGGCCTTCGACGGAAGTTGCAGTGGGATTCAACATTTCAGCGCAATGTTGCGGGACGAGGTCGGTGGCAGGGCCGTTAACCTGCTCCCTTCCGAAACGGTGCAGGACATCTACGGAATCGTGGCAAAAAAGGTAAACGAGATCCTGCAAGCGGATGCCATCAACGGGACGGACAATGAGGTCGTTACGGTGACAGACGAAAATACTGGGGAAATAAGCGAAAAGGTCAAGCTGGGGACCAAAGCACTCGCGGGTCAGTGGCTCGCCTACGGGGTGACACGCTCCGTCACCAAGAGAAGCGTGATGACCCTCGCGTACGGTTCAAAAGAATTCGCCTTCCGCCAGCAAGTGCTGGAGGACACCATCCAGCCGGCGATTGACTCCGGGAAGGGTCTCATGTTTACCCAGCCGAACCAGGCCGCAGGGTACATGGCCAAACTGATCTGGGAAAGCGTTAGCGTCACAGTGGTCGCCGCGGTTGAGGCGATGAATTGGCTGAAGAGCGCGGCAAAGCTCCTCGCCGCTGAGGTGAAGGACAAAAAGACCGGCGAAATCCTGCGCAAGCGCTGCGCCGTCCACTGGGTCACGCCGGATGGATTCCCCGTCTGGCAGGAGTACAAGAAGCCCATCCGAACCCGGCTCAACTTGATGTTCCTTGGCCAGTTTCGCCTGCAGCCCACGATAAACACCAACAAAGACAGCGAGATCGACGCCCACAAGCAGGAGAGCGGCATCGCGCCCAACTTCGTGCACAGTCAGGACGGGTCCCATCTGCGGAAAACTGTTGTGTGGGCTCACGAGAAGTACGGCATTGAGAGCTTCGCCCTGATACACGACAGCTTCGGGACCATACCAGCGGACGCAGCGAACCTGTTCAAAGCCGTGCGGGAAACAATGGTCGACACCTACGAAAGCTGCGACGTACTGGCAGACTTCTATGACCAATTCGCCGACCAGCTTCACGAGTCACAGCTCGACAAGATGCCCGCTCTGCCCGCGAAAGGCAACCTGAATTTGCGCGACATCCTTGAGAGCGATTTTGCGTTCGCCTCTGGTGGTTCTACTAATCTGTCAGATATTATTGAAAAGGAGACCGGTAAGCAACTGGTTATCCAGGAATCCATCCTCATGCTCCCAGAGGAGGTGGAAGAAGTCATTGGGAACAAGCCGGAAAGCGATATACTCGTGCACACCGCCTACGACGAGAGCACCGACGAGAATGTCATGCTTCTGACTAGCGACGCCCCTGAATACAAGCCTTGGGCTCTGGTCATACAGGATAGCAACGGTGAGAACAAGATTAAGATGCTCTCTGGTGGTTCTCCCAAGAAGAAGAGGAAAGTCATGCATTAGACCCAGCTTTCTTGTACAAAGTGGTGATATCCCGCGGCCATGCTAGAGTCCGCAAAAATCACCAGTCTCTCTCTACAAATCTATCTCTCTCTATTTTTCTCCAGAATAATGTGTGAGTAGTTCCCAGATAAGGGAATTAGGGTTCTTATAGGGTTTCGCTCATGTGTTGAGCATATAAGAAACCCTTAGTATGTATTTGTATTTGTAAAATACTTCTATCAATAAAATTTCTAATTCCTAAAACCAAAATCCAGTGACCTGCAGGCATGCGACGTCGGGCCCAAGCTTAGCTTGAGCTTGGATCAGATTGTCGTTTCCCGCCTTCAGTTTAAACTATCAGTGTTTGACAGGATATATTGGCGGGTAAACCTAAGAGAAAAGAGCGTTTATTAGAATAACGGATATTTAAAAGGGCGTGAAAAGGTTTATCCGTTCGTCCATTTGTATGTGCATGCCAACCACAGGGTTCCCCTCGGGATCAAAGTACTTTGATCCAACCCCTCCGCTGCTATAGTGCAGTCGGCTTCTGACGTTCAGTGCAGCCGTCTTCTGAAAACGACATGTCGCACAAGTCCTAAGTTACGCGACAGGCTGCCGCCCTGCCCTTTTCCTGGCGTTTTCTTGTCGCGTGTTTTAGTCGCATAAAGTAGAATACTTGCGACTAGAACCGGAGACATTACGCCATGAACAAGAGCGCCGCCGCTGGCCTGCTGGGCTATGCCCGCGTCAGCACCGACGACCAGGACTTGACCAACCAACGGGCCGAACTGCACGCGGCCGGCTGCACCAAGCTGTTTTCCGAGAAGATCACCGGCACCAGGCGCGACCGCCCGGAGCTGGCCAGGATGCTTGACCACCTACGCCCTGGCGACGTTGTGACAGTGACCAGGCTAGACCGCCTGGCCCGCAGCACCCGCGACCTACTGGACATTGCCGAGCGCATCCAGGAGGCCGGCGCGGGCCTGCGTAGCCTGGCAGAGCCGTGGGCCGACACCACCACGCCGGCCGGCCGCATGGTGTTGACCGTGTTCGCCGGCATTGCCGAGTTCGAGCGTTCCCTAATCATCGACCGCACCCGGAGCGGGCGCGAGGCCGCCAAGGCCCGAGGCGTGAAGTTTGGCCCCCGCCCTACCCTCACCCCGGCACAGATCGCGCACGCCCGCGAGCTGATCGACCAGGAAGGCCGCACCGTGAAAGAGGCGGCTGCACTGCTTGGCGTGCATCGCTCGACCCTGTACCGCGCACTTGAGCGCAGCGAGGAAGTGACGCCCACCGAGGCCAGGCGGCGCGGTGCCTTCCGTGAGGACGCATTGACCGAGGCCGACGCCCTGGCGGCCGCCGAGAATGAACGCCAAGAGGAACAAGCATGAAACCGCACCAGGACGGCCAGGACGAACCGTTTTTCATTACCGAAGAGATCGAGGCGGAGATGATCGCGGCCGGGTACGTGTTCGAGCCGCCCGCGCACGTCTCAACCGTGCGGCTGCATGAAATCCTGGCCGGTTTGTCTGATGCCAAGCTGGCGGCCTGGCCGGCCAGCTTGGCCGCTGAAGAAACCGAGCGCCGCCGTCTAAAAAGGTGATGTGTATTTGAGTAAAACAGCTTGCGTCATGCGGTCGCTGCGTATATGATGCGATGAGTAAATAAACAAATACGCAAGGGGAACGCATGAAGGTTATCGCTGTACTTAACCAGAAAGGCGGGTCAGGCAAGACGACCATCGCAACCCATCTAGCCCGCGCCCTGCAACTCGCCGGGGCCGATGTTCTGTTAGTCGATTCCGATCCCCAGGGCAGTGCCCGCGATTGGGCGGCCGTGCGGGAAGATCAACCGCTAACCGTTGTCGGCATCGACCGCCCGACGATTGACCGCGACGTGAAGGCCATCGGCCGGCGCGACTTCGTAGTGATCGACGGAGCGCCCCAGGCGGCGGACTTGGCTGTGTCCGCGATCAAGGCAGCCGACTTCGTGCTGATTCCGGTGCAGCCAAGCCCTTACGACATATGGGCCACCGCCGACCTGGTGGAGCTGGTTAAGCAGCGCATTGAGGTCACGGATGGAAGGCTACAAGCGGCCTTTGTCGTGTCGCGGGCGATCAAAGGCACGCGCATCGGCGGTGAGGTTGCCGAGGCGCTGGCCGGGTACGAGCTGCCCATTCTTGAGTCCCGTATCACGCAGCGCGTGAGCTACCCAGGCACTGCCGCCGCCGGCACAACCGTTCTTGAATCAGAACCCGAGGGCGACGCTGCCCGCGAGGTCCAGGCGCTGGCCGCTGAAATTAAATCAAAACTCATTTGAGTTAATGAGGTAAAGAGAAAATGAGCAAAAGCACAAACACGCTAAGTGCCGGCCGTCCGAGCGCACGCAGCAGCAAGGCTGCAACGTTGGCCAGCCTGGCAGACACGCCAGCCATGAAGCGGGTCAACTTTCAGTTGCCGGCGGAGGATCACACCAAGCTGAAGATGTACGCGGTACGCCAAGGCAAGACCATTACCGAGCTGCTATCTGAATACATCGCGCAGCTACCAGAGTAAATGAGCAAATGAATAAATGAGTAGATGAATTTTAGCGGCTAAAGGAGGCGGCATGGAAAATCAAGAACAACCAGGCACCGACGCCGTGGAATGCCCCATGTGTGGAGGAACGGGCGGTTGGCCAGGCGTAAGCGGCTGGGTTGTCTGCCGGCCCTGCAATGGCACTGGAACCCCCAAGCCCGAGGAATCGGCGTGACGGTCGCAAACCATCCGGCCCGGTACAAATCGGCGCGGCGCTGGGTGATGACCTGGTGGAGAAGTTGAAGGCCGCGCAGGCCGCCCAGCGGCAACGCATCGAGGCAGAAGCACGCCCCGGTGAATCGTGGCAAGCGGCCGCTGATCGAATCCGCAAAGAATCCCGGCAACCGCCGGCAGCCGGTGCGCCGTCGATTAGGAAGCCGCCCAAGGGCGACGAGCAACCAGATTTTTTCGTTCCGATGCTCTATGACGTGGGCACCCGCGATAGTCGCAGCATCATGGACGTGGCCGTTTTCCGTCTGTCGAAGCGTGACCGACGAGCTGGCGAGGTGATCCGCTACGAGCTTCCAGACGGGCACGTAGAGGTTTCCGCAGGGCCGGCCGGCATGGCCAGTGTGTGGGATTACGACCTGGTACTGATGGCGGTTTCCCATCTAACCGAATCCATGAACCGATACCGGGAAGGGAAGGGAGACAAGCCCGGCCGCGTGTTCCGTCCACACGTTGCGGACGTACTCAAGTTCTGCCGGCGAGCCGATGGCGGAAAGCAGAAAGACGACCTGGTAGAAACCTGCATTCGGTTAAACACCACGCACGTTGCCATGCAGCGTACGAAGAAGGCCAAGAACGGCCGCCTGGTGACGGTATCCGAGGGTGAAGCCTTGATTAGCCGCTACAAGATCGTAAAGAGCGAAACCGGGCGGCCGGAGTACATCGAGATCGAGCTAGCTGATTGGATGTACCGCGAGATCACAGAAGGCAAGAACCCGGACGTGCTGACGGTTCACCCCGATTACTTTTTGATCGATCCCGGCATCGGCCGTTTTCTCTACCGCCTGGCACGCCGCGCCGCAGGCAAGGCAGAAGCCAGATGGTTGTTCAAGACGATCTACGAACGCAGTGGCAGCGCCGGAGAGTTCAAGAAGTTCTGTTTCACCGTGCGCAAGCTGATCGGGTCAAATGACCTGCCGGAGTACGATTTGAAGGAGGAGGCGGGGCAGGCTGGCCCGATCCTAGTCATGCGCTACCGCAACCTGATCGAGGGCGAAGCATCCGCCGGTTCCTAATGTACGGAGCAGATGCTAGGGCAAATTGCCCTAGCAGGGGAAAAAGGTCGAAAAGGTCTCTTTCCTGTGGATAGCACGTACATTGGGAACCCAAAGCCGTACATTGGGAACCGGAACCCGTACATTGGGAACCCAAAGCCGTACATTGGGAACCGGTCACACATGTAAGTGACTGATATAAAAGAGAAAAAAGGCGATTTTTCCGCCTAAAACTCTTTAAAACTTATTAAAACTCTTAAAACCCGCCTGGCCTGTGCATAACTGTCTGGCCAGCGCACAGCCGAAGAGCTGCAAAAAGCGCCTACCCTTCGGTCGCTGCGCTCCCTACGCCCCGCCGCTTCGCGTCGGCCTATCGCGGCCGCTGGCCGCTCAAAAATGGCTGGCCTACGGCCAGGCAATCTACCAGGGCGCGGACAAGCCGCGCCGTCGCCACTCGACCGCCGGCGCCCACATCAAGGCACCCTGCCTCGCGCGTTTCGGTGATGACGGTGAAAACCTCTGACACATGCAGCTCCCGGAGACGGTCACAGCTTGTCTGTAAGCGGATGCCGGGAGCAGACAAGCCCGTCAGGGCGCGTCAGCGGGTGTTGGCGGGTGTCGGGGCGCAGCCATGACCCAGTCACGTAGCGATAGCGGAGTGTATACTGGCTTAACTATGCGGCATCAGAGCAGATTGTACTGAGAGTGCACCATATGCGGTGTGAAATACCGCACAGATGCGTAAGGAGAAAATACCGCATCAGGCGCTCTTCCGCTTCCTCGCTCACTGACTCGCTGCGCTCGGTCGTTCGGCTGCGGCGAGCGGTATCAGCTCACTCAAAGGCGGTAATACGGTTATCCACAGAATCAGGGGATAACGCAGGAAAGAACATGTGAGCAAAAGGCCAGCAAAAGGCCAGGAACCGTAAAAAGGCCGCGTTGCTGGCGTTTTTCCATAGGCTCCGCCCCCCTGACGAGCATCACAAAAATCGACGCTCAAGTCAGAGGTGGCGAAACCCGACAGGACTATAAAGATACCAGGCGTTTCCCCCTGGAAGCTCCCTCGTGCGCTCTCCTGTTCCGACCCTGCCGCTTACCGGATACCTGTCCGCCTTTCTCCCTTCGGGAAGCGTGGCGCTTTCTCATAGCTCACGCTGTAGGTATCTCAGTTCGGTGTAGGTCGTTCGCTCCAAGCTGGGCTGTGTGCACGAACCCCCCGTTCAGCCCGACCGCTGCGCCTTATCCGGTAACTATCGTCTTGAGTCCAACCCGGTAAGACACGACTTATCGCCACTGGCAGCAGCCACTGGTAACAGGATTAGCAGAGCGAGGTATGTAGGCGGTGCTACAGAGTTCTTGAAGTGGTGGCCTAACTACGGCTACACTAGAAGGACAGTATTTGGTATCTGCGCTCTGCTGAAGCCAGTTACCTTCGGAAAAAGAGTTGGTAGCTCTTGATCCGGCAAACAAACCACCGCTGGTAGCGGTGGTTTTTTTGTTTGCAAGCAGCAGATTACGCGCAGAAAAAAAGGATCTCAAGAAGATCCTTTGATCTTTTCTACGGGGTCTGACGCTCAGTGGAACGAAAACTCACGTTAAGGGATTTTGGTCATGCATGATATATCTCCCAATTTGTGTAGGGCTTATTATGCACGCTTAAAAATAATAAAAGCAGACTTGACCTGATAGTTTGGCTGTGAGCAATTATGTGCTTAGTGCATCTAATCGCTTGAGTTAACGCCGGCGAAGCGGCGTCGGCTTGAACGAATTTCTAGCTAGACATTATTTGCCGACTACCTTGGTGATCTCGCCTTTCACGTAGTGGACAAATTCTTCCAACTGATCTGCGCGCGAGGCCAAGCGATCTTCTTCTTGTCCAAGATAAGCCTGTCTAGCTTCAAGTATGACGGGCTGATACTGGGCCGGCAGGCGCTCCATTGCCCAGTCGGCAGCGACATCCTTCGGCGCGATTTTGCCGGTTACTGCGCTGTACCAAATGCGGGACAACGTAAGCACTACATTTCGCTCATCGCCAGCCCAGTCGGGCGGCGAGTTCCATAGCGTTAAGGTTTCATTTAGCGCCTCAAATAGATCCTGTTCAGGAACCGGATCAAAGAGTTCCTCCGCCGCTGGACCTACCAAGGCAACGCTATGTTCTCTTGCTTTTGTCAGCAAGATAGCCAGATCAATGTCGATCGTGGCTGGCTCGAAGATACCTGCAAGAATGTCATTGCGCTGCCATTCTCCAAATTGCAGTTCGCGCTTAGCTGGATAACGCCACGGAATGATGTCGTCGTGCACAACAATGGTGACTTCTACAGCGCGGAGAATCTCGCTCTCTCCAGGGGAAGCCGAAGTTTCCAAAAGGTCGTTGATCAAAGCTCGCCGCGTTGTTTCATCAAGCCTTACGGTCACCGTAACCAGCAAATCAATATCACTGTGTGGCTTCAGGCCGCCATCCACTGCGGAGCCGTACAAATGTACGGCCAGCAACGTCGGTTCGAGATGGCGCTCGATGACGCCAACTACCTCTGATAGTTGAGTCGATACTTCGGCGATCACCGCTTCCCCCATGATGTTTAACTTTGTTTTAGGGCGACTGCCCTGCTGCGTAACATCGTTGCTGCTCCATAACATCAAACATCGACCCACGGCGTAACGCGCTTGCTGCTTGGATGCCCGAGGCATAGACTGTACCCCAAAAAAACATGTCATAACAAGAAGCCATGAAAACCGCCACTGCGCCGTTACCACCGCTGCGTTCGGTCAAGGTTCTGGACCAGTTGCGTGACGGCAGTTACGCTACTTGCATTACAGCTTACGAACCGAACGAGGCTTATGTCCACTGGGTTCGTGCCCGAATTGATCACAGGCAGCAACGCTCTGTCATCGTTACAATCAACATGCTACCCTCCGCGAGATCATCCGTGTTTCAAACCCGGCAGCTTAGTTGCCGTTCTTCCGAATAGCATCGGTAACATGAGCAAAGTCTGCCGCCTTACAACGGCTCTCCCGCTGACGCCGTCCCGGACTGATGGGCTGCCTGTATCGAGTGGTGATTTTGTGCCGAGCTGCCGGTCGGGGAGCTGTTGGCTGGCTGGTGGCAGGATATATTGTGGTGTAAACAAATTGACGCTTAGACAACTTAATAACACATTGCGGACGTTTTTAATGTACTGAATTAACGCCGAATTGAATTATCAGCTTGCATGCCGGTCGATCTAGTAACATAGATGACACCGCGCGCGATAATTTATCCTAGTTTGCGCGCTATATTTTGTTTTCTATCGCGTATTAAATGTATAATTGCGGGACTCTAATCATAAAAACCCATCTCATAAATAACGTCATGCATTACATGTTAATTATTACATGCTTAACGTAATTCAACAGAAATTATATGATAATCATCGCAAGACCGGCAACAGGATTCAATCTTAAGAAACTTTATTGCCAAATGTTTGAACGATCTGCTTGACTCTAGCTAGAGTCCGAACCCCAGAGTCCCGCTCAGAAGAACTCGTCAAGAAGGCGATAGAAGGCGATGCGCTGCGAATCGGGAGCGGCGATACCGTAAAGCACGAGGAAGCGGTCAGCCCATTCGCCGCCAAGCTCTTCAGCAATATCACGGGTAGCCAACGCTATGTCCTGATAGCGGTCCGCCACACCCAGCCGGCCACAGTCGATGAATCCAGAAAAGCGGCCATTTTCCACCATGATATTCGGCAAGCAGGCATCGCCGTGGGTCACGACGAGATCCTCGCCGTCGGGCATCCGCGCCTTGAGCCTGGCGAACAGTTCGGCTGGCGCGAGCCCCTGATGCTCTTCGTCCAGATCATCCTGATCGACAAGACCGGCTTCCATCCGAGTACGTGCTCGCTCGATGCGATGTTTCGCTTGGTGGTCGAATGGGCAGGTAGCCGGATCAAGCGTATGCAGCCGCCGCATTGCATCAGCCATGATGGATACTTTCTCGGCAGGAGCAAGGTGAGATGACAGGAGATCCTGCCCCGGCACTTCGCCCAATAGCAGCCAGTCCCTTCCCGCTTCAGTGACAACGTCGAGCACAGCTGCGCAAGGAACGCCCGTCGTGGCCAGCCACGATAGCCGCGCTGCCTCGTCTTGGAGTTCATTCAGGGCACCGGACAGGTCGGTCTTGACAAAAAGAACCGGGCGCCCCTGCGCTGACAGCCGGAACACGGCGGCATCAGAGCAGCCGATTGTCTGTTGTGCCCAGTCATAGCCGAATAGCCTCTCCACCCAAGCGGCCGGAGAACCTGCGTGCAATCCATCTTGTTCAATCATGCCTCGATCGAGTTGAGAGTGAATATGAGACTCTAATTGGATACCGAGGGGAATTTATGGAACGTCAGTGGAGCATTTTTGACAAGAAATATTTGCTAGCTGATAGTGACCTTAGGCGACTTTTGAACGCGCAATAATGGTTTCTGACGTATGTGCTTAGCTCATTAAACTCCAGAAACCCGCGGCTGAGTGGCTCCTTCAACGTTGCGGTTCTGTCAGTTCCAAACGTAAAACGGCTTGTCCCGCGTCATCGGCGGGGGTCATAACGTGACTCCCTTAATTCTCATGTATGATAATTCGAGCT

**Sequences S3:** DNA sequence of the vector pEditor: **p35S-AID-T7pol** (p35S_3xFLAG_AID_T7_UGI _pK2GW7) (two-vector system)

CTCCCATATGGTCGACTAGAGCCAAGCTGATCTCCTTTGCCCCGGAGATCACCATGGACGACTTTCTCTATCTCTACGATCTAGGAAGAAAGTTCGACGGAGAAGGTGACGATACCATGTTCACCACCGATAATGAGAAGATTAGCCTCTTCAATTTCAGAAAGAATGCTGACCCACAGATGGTTAGAGAGGCCTACGCGGCAGGTCTCATCAAGACGATCTACCCGAGTAATAATCTCCAGGAGATCAAATACCTTCCCAAGAAGGTTAAAGATGCAGTCAAAAGATTCAGGACTAACTGCATCAAGAACACAGAGAAAGATATATTTCTCAAGATCAGAAGTACTATTCCAGTATGGACGATTCAAGGCTTGCTTCATAAACCAAGGCAAGTAATAGAGATTGGAGTCTCTAAGAAAGTAGTTCCTACTGAATCAAAGGCCATGGAGTCAAAAATTCAGATCGAGGATCTAACAGAACTCGCCGTGAAGACTGGCGAACAGTTCATACAGAGTCTTTTACGACTCAATGACAAGAAGAAAATCTTCGTCAACATGGTGGAGCACGACACTCTCGTCTACTCCAAGAATATCAAAGATACAGTCTCAGAAGACCAAAGGGCTATTGAGACTTTTCAACAAAGGGTAATATCGGGAAACCTCCTCGGATTCCATTGCCCAGCTATCTGTCACTTCATCAAAAGGACAGTAGAAAAGGAAGGTGGCACCTACAAATGCCATCATTGCGATAAAGGAAAGGCTATCGTTCAAGATGCCTCTGCCGACAGTGGTCCCAAAGATGGACCCCCACCCACGAGGAGCATCGTGGAAAAAGAAGACGTTCCAACCACGTCTTCAAAGCAAGTGGATTGATGTGATATCTCCACTGACGTAAGGGATGACGCACAATCCCACTATCCTTCGCAAGACCCTTCCTCTATATAAGGAAGTTCATTTCATTTGGAGAGGACTCCGGTATTTTTACAACAATACCACAACAAAACAAACAACAAACAACATTACAATTTACTATTCTAGTCGACCTGCAGGCGGCCGCACTAGTGATATCACAAGTTTGTACAAAAAAGCAGGCTTCGAAGGAGATAGAACCAATTCTCTAAGGAAATACTTAACCATGGACTATAAGGACCACGACGGAGACTACAAGGATCATGATATTGATTACAAAGACGATGACGATAAGATGGCCCCAAAGAAGAAGCGGAAGGTGGGTATCCACGGAGTCCCAGCAGCCGCATGCCCCGGGATGGACAGCCTCTTGATGAACCGGAGGGAGTTTCTTTACCAATTCAAAAATGTCCGCTGGGCTAAGGGTCGGCGTGAGACCTACCTGTGCTACGTAGTGAAGAGGCGTGACAGTGCTACATCCTTTTCACTGGACTTTGGTTATCTTCGCAATAAGAACGGCTGCCACGTGGAATTGCTCTTCCTCCGCTACATCTCGGACTGGGACCTAGACCCTGGCCGCTGCTACCGCGTCACCTGGTTCATCTCCTGGAGCCCCTGCTACGACTGTGCCCGACATGTGGCCGACTTTCTGCGAGGGAACCCCAACCTCAGTCTGAGGATCTTCACCGCGCGCCTCTACTTCTGTGAGGACCGCAAGGCTGAGCCCGAGGGGCTGCGGCGGCTGCACCGCGCCGGGGTGCAAATAGCCATCATGACCTTCAAAGATTATTTTTACTGCTGGAATACTTTTGTAGAAAACCACGGAAGAACTTTCAAAGCCTGGGAAGGGCTGCATGAAAATTCAGTTCGTCTCTCCAGACAGCTTCGGCGCATCCTTTTGCCCCTGTATGAGGTTGATGACTTACGAGACGCATTTCGTACTAGCGGCAGCGAGACTCCCGGGACCTCAGAGTCCGCCACACCCGAAAGTAACACCATCAACATTGCTAAGAACGACTTCTCAGACATAGAGCTCGCGGCTATTCCGTTCAACACCCTGGCTGACCACTACGGCGAGAGACTCGCTAGGGAGCAGCTGGCGTTGGAGCATGAATCCTACGAGATGGGCGAGGCTAGGTTCCGCAAGATGTTCGAGCGACAATTGAAGGCAGGGGAGGTGGCGGACAACGCTGCCGCCAAGCCCCTGATCACAACCTTGCTGCCCAAAATGATCGCGCGGATCAACGATTGGTTTGAGGAGGTTAAGGCAAAACGGGGCAAACGCCCGACCGCATTTCAATTCCTCCAAGAAATCAAGCCTGAGGCTGTTGCCTACATCACTATCAAGACGACACTGGCGTGTCTCACAAGCGCCGACAACACCACCGTGCAAGCCGTCGCCAGCGCCATCGGGCGGGCAATTGAGGATGAGGCACGGTTTGGTAGGATCCGAGACCTGGAAGCGAAGCACTTCAAGAAGAACGTGGAAGAGCAGTTGAACAAACGCGTCGGCCACGTGTATAAAAAGGCTTTCATGCAGGTGGTGGAGGCCGATATGCTCAGTAAGGGGCTGCTTGGGGGGGAGGCGTGGTCATCCTGGCACAAGGAGGATAGCATTCACGTGGGGGTCCGATGTATCGAGATGCTGATAGAGAGCACCGGAATGGTCTCCCTCCATCGCCAGAACGCTGGGGTCGTAGGGCAGGACTCCGAGACTATTGAGCTGGCCCCCGAGTATGCCGAAGCAATCGCTACACGCGCAGGTGCACTGGCTGGGATAAGCCCTATGTTTCAGCCCTGCGTAGTGCCTCCAAAGCCATGGACCGGCATCACAGGGGGTGGCTATTGGGCCAACGGTAGGCGGCCTCTGGCCCTGGTACGCACGCACAGCAAGAAGGCGCTCATGCGCTATGAAGACGTTTACATGCCCGAGGTTTACAAGGCGATCAATATCGCGCAGAACACCGCCTGGAAAATCAATAAGAAGGTGTTGGCGGTCGCAAACGTGATTACCAAGTGGAAGCATTGCCCAGTCGAGGACATACCCGCCATAGAACGCGAAGAGCTGCCGATGAAGCCGGAAGACATTGATATGAACCCCGAGGCCCTCACCGCGTGGAAAAGAGCCGCAGCCGCCGTATACAGGAAGGATAAAGCGCGCAAGTCCCTACGCATAAGCCTCGAGTTTATGCTGGAACAGGCCAACAAGTTCGCCAACCACAAAGCTATCTGGTTCCCCTACAACATGGACTGGAGAGGGAGGGTCTACGCCGTCAGCATGTTCAATCCCCAGGGCAACGACATGACGAAGGGCCTTCTGACATTGGCAAAGGGGAAGCCTATCGGAAAGGAGGGGTACTACTGGCTCAAGATCCACGGCGCCAACTGCGCGGGAGTGGACAAGGTTCCATTTCCCGAGCGAATTAAGTTCATCGAGGAAAACCACGAAAACATTATGGCGTGCGCTAAATCCCCCCTCGAGAACACATGGTGGGCCGAGCAAGACTCCCCGTTCTGTTTTTTGGCATTCTGCTTTGAGTACGCCGGTGTGCAGCACCATGGCCTCTCATACAACTGTTCCCTGCCCCTGGCCTTCGACGGAAGTTGCAGTGGGATTCAACATTTCAGCGCAATGTTGCGGGACGAGGTCGGTGGCAGGGCCGTTAACCTGCTCCCTTCCGAAACGGTGCAGGACATCTACGGAATCGTGGCAAAAAAGGTAAACGAGATCCTGCAAGCGGATGCCATCAACGGGACGGACAATGAGGTCGTTACGGTGACAGACGAAAATACTGGGGAAATAAGCGAAAAGGTCAAGCTGGGGACCAAAGCACTCGCGGGTCAGTGGCTCGCCTACGGGGTGACACGCTCCGTCACCAAGAGAAGCGTGATGACCCTCGCGTACGGTTCAAAAGAATTCGCCTTCCGCCAGCAAGTGCTGGAGGACACCATCCAGCCGGCGATTGACTCCGGGAAGGGTCTCATGTTTACCCAGCCGAACCAGGCCGCAGGGTACATGGCCAAACTGATCTGGGAAAGCGTTAGCGTCACAGTGGTCGCCGCGGTTGAGGCGATGAATTGGCTGAAGAGCGCGGCAAAGCTCCTCGCCGCTGAGGTGAAGGACAAAAAGACCGGCGAAATCCTGCGCAAGCGCTGCGCCGTCCACTGGGTCACGCCGGATGGATTCCCCGTCTGGCAGGAGTACAAGAAGCCCATCCGAACCCGGCTCAACTTGATGTTCCTTGGCCAGTTTCGCCTGCAGCCCACGATAAACACCAACAAAGACAGCGAGATCGACGCCCACAAGCAGGAGAGCGGCATCGCGCCCAACTTCGTGCACAGTCAGGACGGGTCCCATCTGCGGAAAACTGTTGTGTGGGCTCACGAGAAGTACGGCATTGAGAGCTTCGCCCTGATACACGACAGCTTCGGGACCATACCAGCGGACGCAGCGAACCTGTTCAAAGCCGTGCGGGAAACAATGGTCGACACCTACGAAAGCTGCGACGTACTGGCAGACTTCTATGACCAATTCGCCGACCAGCTTCACGAGTCACAGCTCGACAAGATGCCCGCTCTGCCCGCGAAAGGCAACCTGAATTTGCGCGACATCCTTGAGAGCGATTTTGCGTTCGCCTCTGGTGGTTCTACTAATCTGTCAGATATTATTGAAAAGGAGACCGGTAAGCAACTGGTTATCCAGGAATCCATCCTCATGCTCCCAGAGGAGGTGGAAGAAGTCATTGGGAACAAGCCGGAAAGCGATATACTCGTGCACACCGCCTACGACGAGAGCACCGACGAGAATGTCATGCTTCTGACTAGCGACGCCCCTGAATACAAGCCTTGGGCTCTGGTCATACAGGATAGCAACGGTGAGAACAAGATTAAGATGCTCTCTGGTGGTTCTCCCAAGAAGAAGAGGAAAGTCATGCATTAGACCCAGCTTTCTTGTACAAAGTGGTGATATCCCGCGGCCATGCTAGAGTCCGCAAAAATCACCAGTCTCTCTCTACAAATCTATCTCTCTCTATTTTTCTCCAGAATAATGTGTGAGTAGTTCCCAGATAAGGGAATTAGGGTTCTTATAGGGTTTCGCTCATGTGTTGAGCATATAAGAAACCCTTAGTATGTATTTGTATTTGTAAAATACTTCTATCAATAAAATTTCTAATTCCTAAAACCAAAATCCAGTGACCTGCAGGCATGCGACGTCGGGCCCAAGCTTAGCTTGAGCTTGGATCAGATTGTCGTTTCCCGCCTTCAGTTTAAACTATCAGTGTTTGACAGGATATATTGGCGGGTAAACCTAAGAGAAAAGAGCGTTTATTAGAATAACGGATATTTAAAAGGGCGTGAAAAGGTTTATCCGTTCGTCCATTTGTATGTGCATGCCAACCACAGGGTTCCCCTCGGGATCAAAGTACTTTGATCCAACCCCTCCGCTGCTATAGTGCAGTCGGCTTCTGACGTTCAGTGCAGCCGTCTTCTGAAAACGACATGTCGCACAAGTCCTAAGTTACGCGACAGGCTGCCGCCCTGCCCTTTTCCTGGCGTTTTCTTGTCGCGTGTTTTAGTCGCATAAAGTAGAATACTTGCGACTAGAACCGGAGACATTACGCCATGAACAAGAGCGCCGCCGCTGGCCTGCTGGGCTATGCCCGCGTCAGCACCGACGACCAGGACTTGACCAACCAACGGGCCGAACTGCACGCGGCCGGCTGCACCAAGCTGTTTTCCGAGAAGATCACCGGCACCAGGCGCGACCGCCCGGAGCTGGCCAGGATGCTTGACCACCTACGCCCTGGCGACGTTGTGACAGTGACCAGGCTAGACCGCCTGGCCCGCAGCACCCGCGACCTACTGGACATTGCCGAGCGCATCCAGGAGGCCGGCGCGGGCCTGCGTAGCCTGGCAGAGCCGTGGGCCGACACCACCACGCCGGCCGGCCGCATGGTGTTGACCGTGTTCGCCGGCATTGCCGAGTTCGAGCGTTCCCTAATCATCGACCGCACCCGGAGCGGGCGCGAGGCCGCCAAGGCCCGAGGCGTGAAGTTTGGCCCCCGCCCTACCCTCACCCCGGCACAGATCGCGCACGCCCGCGAGCTGATCGACCAGGAAGGCCGCACCGTGAAAGAGGCGGCTGCACTGCTTGGCGTGCATCGCTCGACCCTGTACCGCGCACTTGAGCGCAGCGAGGAAGTGACGCCCACCGAGGCCAGGCGGCGCGGTGCCTTCCGTGAGGACGCATTGACCGAGGCCGACGCCCTGGCGGCCGCCGAGAATGAACGCCAAGAGGAACAAGCATGAAACCGCACCAGGACGGCCAGGACGAACCGTTTTTCATTACCGAAGAGATCGAGGCGGAGATGATCGCGGCCGGGTACGTGTTCGAGCCGCCCGCGCACGTCTCAACCGTGCGGCTGCATGAAATCCTGGCCGGTTTGTCTGATGCCAAGCTGGCGGCCTGGCCGGCCAGCTTGGCCGCTGAAGAAACCGAGCGCCGCCGTCTAAAAAGGTGATGTGTATTTGAGTAAAACAGCTTGCGTCATGCGGTCGCTGCGTATATGATGCGATGAGTAAATAAACAAATACGCAAGGGGAACGCATGAAGGTTATCGCTGTACTTAACCAGAAAGGCGGGTCAGGCAAGACGACCATCGCAACCCATCTAGCCCGCGCCCTGCAACTCGCCGGGGCCGATGTTCTGTTAGTCGATTCCGATCCCCAGGGCAGTGCCCGCGATTGGGCGGCCGTGCGGGAAGATCAACCGCTAACCGTTGTCGGCATCGACCGCCCGACGATTGACCGCGACGTGAAGGCCATCGGCCGGCGCGACTTCGTAGTGATCGACGGAGCGCCCCAGGCGGCGGACTTGGCTGTGTCCGCGATCAAGGCAGCCGACTTCGTGCTGATTCCGGTGCAGCCAAGCCCTTACGACATATGGGCCACCGCCGACCTGGTGGAGCTGGTTAAGCAGCGCATTGAGGTCACGGATGGAAGGCTACAAGCGGCCTTTGTCGTGTCGCGGGCGATCAAAGGCACGCGCATCGGCGGTGAGGTTGCCGAGGCGCTGGCCGGGTACGAGCTGCCCATTCTTGAGTCCCGTATCACGCAGCGCGTGAGCTACCCAGGCACTGCCGCCGCCGGCACAACCGTTCTTGAATCAGAACCCGAGGGCGACGCTGCCCGCGAGGTCCAGGCGCTGGCCGCTGAAATTAAATCAAAACTCATTTGAGTTAATGAGGTAAAGAGAAAATGAGCAAAAGCACAAACACGCTAAGTGCCGGCCGTCCGAGCGCACGCAGCAGCAAGGCTGCAACGTTGGCCAGCCTGGCAGACACGCCAGCCATGAAGCGGGTCAACTTTCAGTTGCCGGCGGAGGATCACACCAAGCTGAAGATGTACGCGGTACGCCAAGGCAAGACCATTACCGAGCTGCTATCTGAATACATCGCGCAGCTACCAGAGTAAATGAGCAAATGAATAAATGAGTAGATGAATTTTAGCGGCTAAAGGAGGCGGCATGGAAAATCAAGAACAACCAGGCACCGACGCCGTGGAATGCCCCATGTGTGGAGGAACGGGCGGTTGGCCAGGCGTAAGCGGCTGGGTTGTCTGCCGGCCCTGCAATGGCACTGGAACCCCCAAGCCCGAGGAATCGGCGTGACGGTCGCAAACCATCCGGCCCGGTACAAATCGGCGCGGCGCTGGGTGATGACCTGGTGGAGAAGTTGAAGGCCGCGCAGGCCGCCCAGCGGCAACGCATCGAGGCAGAAGCACGCCCCGGTGAATCGTGGCAAGCGGCCGCTGATCGAATCCGCAAAGAATCCCGGCAACCGCCGGCAGCCGGTGCGCCGTCGATTAGGAAGCCGCCCAAGGGCGACGAGCAACCAGATTTTTTCGTTCCGATGCTCTATGACGTGGGCACCCGCGATAGTCGCAGCATCATGGACGTGGCCGTTTTCCGTCTGTCGAAGCGTGACCGACGAGCTGGCGAGGTGATCCGCTACGAGCTTCCAGACGGGCACGTAGAGGTTTCCGCAGGGCCGGCCGGCATGGCCAGTGTGTGGGATTACGACCTGGTACTGATGGCGGTTTCCCATCTAACCGAATCCATGAACCGATACCGGGAAGGGAAGGGAGACAAGCCCGGCCGCGTGTTCCGTCCACACGTTGCGGACGTACTCAAGTTCTGCCGGCGAGCCGATGGCGGAAAGCAGAAAGACGACCTGGTAGAAACCTGCATTCGGTTAAACACCACGCACGTTGCCATGCAGCGTACGAAGAAGGCCAAGAACGGCCGCCTGGTGACGGTATCCGAGGGTGAAGCCTTGATTAGCCGCTACAAGATCGTAAAGAGCGAAACCGGGCGGCCGGAGTACATCGAGATCGAGCTAGCTGATTGGATGTACCGCGAGATCACAGAAGGCAAGAACCCGGACGTGCTGACGGTTCACCCCGATTACTTTTTGATCGATCCCGGCATCGGCCGTTTTCTCTACCGCCTGGCACGCCGCGCCGCAGGCAAGGCAGAAGCCAGATGGTTGTTCAAGACGATCTACGAACGCAGTGGCAGCGCCGGAGAGTTCAAGAAGTTCTGTTTCACCGTGCGCAAGCTGATCGGGTCAAATGACCTGCCGGAGTACGATTTGAAGGAGGAGGCGGGGCAGGCTGGCCCGATCCTAGTCATGCGCTACCGCAACCTGATCGAGGGCGAAGCATCCGCCGGTTCCTAATGTACGGAGCAGATGCTAGGGCAAATTGCCCTAGCAGGGGAAAAAGGTCGAAAAGGTCTCTTTCCTGTGGATAGCACGTACATTGGGAACCCAAAGCCGTACATTGGGAACCGGAACCCGTACATTGGGAACCCAAAGCCGTACATTGGGAACCGGTCACACATGTAAGTGACTGATATAAAAGAGAAAAAAGGCGATTTTTCCGCCTAAAACTCTTTAAAACTTATTAAAACTCTTAAAACCCGCCTGGCCTGTGCATAACTGTCTGGCCAGCGCACAGCCGAAGAGCTGCAAAAAGCGCCTACCCTTCGGTCGCTGCGCTCCCTACGCCCCGCCGCTTCGCGTCGGCCTATCGCGGCCGCTGGCCGCTCAAAAATGGCTGGCCTACGGCCAGGCAATCTACCAGGGCGCGGACAAGCCGCGCCGTCGCCACTCGACCGCCGGCGCCCACATCAAGGCACCCTGCCTCGCGCGTTTCGGTGATGACGGTGAAAACCTCTGACACATGCAGCTCCCGGAGACGGTCACAGCTTGTCTGTAAGCGGATGCCGGGAGCAGACAAGCCCGTCAGGGCGCGTCAGCGGGTGTTGGCGGGTGTCGGGGCGCAGCCATGACCCAGTCACGTAGCGATAGCGGAGTGTATACTGGCTTAACTATGCGGCATCAGAGCAGATTGTACTGAGAGTGCACCATATGCGGTGTGAAATACCGCACAGATGCGTAAGGAGAAAATACCGCATCAGGCGCTCTTCCGCTTCCTCGCTCACTGACTCGCTGCGCTCGGTCGTTCGGCTGCGGCGAGCGGTATCAGCTCACTCAAAGGCGGTAATACGGTTATCCACAGAATCAGGGGATAACGCAGGAAAGAACATGTGAGCAAAAGGCCAGCAAAAGGCCAGGAACCGTAAAAAGGCCGCGTTGCTGGCGTTTTTCCATAGGCTCCGCCCCCCTGACGAGCATCACAAAAATCGACGCTCAAGTCAGAGGTGGCGAAACCCGACAGGACTATAAAGATACCAGGCGTTTCCCCCTGGAAGCTCCCTCGTGCGCTCTCCTGTTCCGACCCTGCCGCTTACCGGATACCTGTCCGCCTTTCTCCCTTCGGGAAGCGTGGCGCTTTCTCATAGCTCACGCTGTAGGTATCTCAGTTCGGTGTAGGTCGTTCGCTCCAAGCTGGGCTGTGTGCACGAACCCCCCGTTCAGCCCGACCGCTGCGCCTTATCCGGTAACTATCGTCTTGAGTCCAACCCGGTAAGACACGACTTATCGCCACTGGCAGCAGCCACTGGTAACAGGATTAGCAGAGCGAGGTATGTAGGCGGTGCTACAGAGTTCTTGAAGTGGTGGCCTAACTACGGCTACACTAGAAGGACAGTATTTGGTATCTGCGCTCTGCTGAAGCCAGTTACCTTCGGAAAAAGAGTTGGTAGCTCTTGATCCGGCAAACAAACCACCGCTGGTAGCGGTGGTTTTTTTGTTTGCAAGCAGCAGATTACGCGCAGAAAAAAAGGATCTCAAGAAGATCCTTTGATCTTTTCTACGGGGTCTGACGCTCAGTGGAACGAAAACTCACGTTAAGGGATTTTGGTCATGCATGATATATCTCCCAATTTGTGTAGGGCTTATTATGCACGCTTAAAAATAATAAAAGCAGACTTGACCTGATAGTTTGGCTGTGAGCAATTATGTGCTTAGTGCATCTAATCGCTTGAGTTAACGCCGGCGAAGCGGCGTCGGCTTGAACGAATTTCTAGCTAGACATTATTTGCCGACTACCTTGGTGATCTCGCCTTTCACGTAGTGGACAAATTCTTCCAACTGATCTGCGCGCGAGGCCAAGCGATCTTCTTCTTGTCCAAGATAAGCCTGTCTAGCTTCAAGTATGACGGGCTGATACTGGGCCGGCAGGCGCTCCATTGCCCAGTCGGCAGCGACATCCTTCGGCGCGATTTTGCCGGTTACTGCGCTGTACCAAATGCGGGACAACGTAAGCACTACATTTCGCTCATCGCCAGCCCAGTCGGGCGGCGAGTTCCATAGCGTTAAGGTTTCATTTAGCGCCTCAAATAGATCCTGTTCAGGAACCGGATCAAAGAGTTCCTCCGCCGCTGGACCTACCAAGGCAACGCTATGTTCTCTTGCTTTTGTCAGCAAGATAGCCAGATCAATGTCGATCGTGGCTGGCTCGAAGATACCTGCAAGAATGTCATTGCGCTGCCATTCTCCAAATTGCAGTTCGCGCTTAGCTGGATAACGCCACGGAATGATGTCGTCGTGCACAACAATGGTGACTTCTACAGCGCGGAGAATCTCGCTCTCTCCAGGGGAAGCCGAAGTTTCCAAAAGGTCGTTGATCAAAGCTCGCCGCGTTGTTTCATCAAGCCTTACGGTCACCGTAACCAGCAAATCAATATCACTGTGTGGCTTCAGGCCGCCATCCACTGCGGAGCCGTACAAATGTACGGCCAGCAACGTCGGTTCGAGATGGCGCTCGATGACGCCAACTACCTCTGATAGTTGAGTCGATACTTCGGCGATCACCGCTTCCCCCATGATGTTTAACTTTGTTTTAGGGCGACTGCCCTGCTGCGTAACATCGTTGCTGCTCCATAACATCAAACATCGACCCACGGCGTAACGCGCTTGCTGCTTGGATGCCCGAGGCATAGACTGTACCCCAAAAAAACATGTCATAACAAGAAGCCATGAAAACCGCCACTGCGCCGTTACCACCGCTGCGTTCGGTCAAGGTTCTGGACCAGTTGCGTGACGGCAGTTACGCTACTTGCATTACAGCTTACGAACCGAACGAGGCTTATGTCCACTGGGTTCGTGCCCGAATTGATCACAGGCAGCAACGCTCTGTCATCGTTACAATCAACATGCTACCCTCCGCGAGATCATCCGTGTTTCAAACCCGGCAGCTTAGTTGCCGTTCTTCCGAATAGCATCGGTAACATGAGCAAAGTCTGCCGCCTTACAACGGCTCTCCCGCTGACGCCGTCCCGGACTGATGGGCTGCCTGTATCGAGTGGTGATTTTGTGCCGAGCTGCCGGTCGGGGAGCTGTTGGCTGGCTGGTGGCAGGATATATTGTGGTGTAAACAAATTGACGCTTAGACAACTTAATAACACATTGCGGACGTTTTTAATGTACTGAATTAACGCCGAATTGAATTATCAGCTTGCATGCCGGTCGATCTAGTAACATAGATGACACCGCGCGCGATAATTTATCCTAGTTTGCGCGCTATATTTTGTTTTCTATCGCGTATTAAATGTATAATTGCGGGACTCTAATCATAAAAACCCATCTCATAAATAACGTCATGCATTACATGTTAATTATTACATGCTTAACGTAATTCAACAGAAATTATATGATAATCATCGCAAGACCGGCAACAGGATTCAATCTTAAGAAACTTTATTGCCAAATGTTTGAACGATCTGCTTGACTCTAGCTAGAGTCCGAACCCCAGAGTCCCGCTCAGAAGAACTCGTCAAGAAGGCGATAGAAGGCGATGCGCTGCGAATCGGGAGCGGCGATACCGTAAAGCACGAGGAAGCGGTCAGCCCATTCGCCGCCAAGCTCTTCAGCAATATCACGGGTAGCCAACGCTATGTCCTGATAGCGGTCCGCCACACCCAGCCGGCCACAGTCGATGAATCCAGAAAAGCGGCCATTTTCCACCATGATATTCGGCAAGCAGGCATCGCCGTGGGTCACGACGAGATCCTCGCCGTCGGGCATCCGCGCCTTGAGCCTGGCGAACAGTTCGGCTGGCGCGAGCCCCTGATGCTCTTCGTCCAGATCATCCTGATCGACAAGACCGGCTTCCATCCGAGTACGTGCTCGCTCGATGCGATGTTTCGCTTGGTGGTCGAATGGGCAGGTAGCCGGATCAAGCGTATGCAGCCGCCGCATTGCATCAGCCATGATGGATACTTTCTCGGCAGGAGCAAGGTGAGATGACAGGAGATCCTGCCCCGGCACTTCGCCCAATAGCAGCCAGTCCCTTCCCGCTTCAGTGACAACGTCGAGCACAGCTGCGCAAGGAACGCCCGTCGTGGCCAGCCACGATAGCCGCGCTGCCTCGTCTTGGAGTTCATTCAGGGCACCGGACAGGTCGGTCTTGACAAAAAGAACCGGGCGCCCCTGCGCTGACAGCCGGAACACGGCGGCATCAGAGCAGCCGATTGTCTGTTGTGCCCAGTCATAGCCGAATAGCCTCTCCACCCAAGCGGCCGGAGAACCTGCGTGCAATCCATCTTGTTCAATCATGCCTCGATCGAGTTGAGAGTGAATATGAGACTCTAATTGGATACCGAGGGGAATTTATGGAACGTCAGTGGAGCATTTTTGACAAGAAATATTTGCTAGCTGATAGTGACCTTAGGCGACTTTTGAACGCGCAATAATGGTTTCTGACGTATGTGCTTAGCTCATTAAACTCCAGAAACCCGCGGCTGAGTGGCTCCTTCAACGTTGCGGTTCTGTCAGTTCCAAACGTAAAACGGCTTGTCCCGCGTCATCGGCGGGGGTCATAACGTGACTCCCTTAATTCTCATGTATGATAATTCGAGCT

**Sequences S4:** DNA sequence of the vector pTarget: **p35S-pT7-GFP** (p35S_pT7_GFP_NOSterm) (two-vector system)

GGTAATACGACTCACTATAGGGAGACCACAACGGTTTCCCTAGAAATAATTTTGTTTAACTTTAAGAAGGAGATATATCTAGAATGAAGACTAATCTTTTTCTCTTTCTCATCTTTTCACTTCTCCTATCATTATCCTCGGCCGAATTCAGTAAAGGAGAAGAACTTTTCACTGGAGTTGTCCCAATTCTTGTTGAATTAGATGGTGATGTTAATGGGCACAAATTTTCTGTCAGTGGAGAGGGTGAAGGTGATGCAACATACGGAAAACTTACCCTTAAATTTATTTGCACTACTGGAAAACTACCTGTTCCATGGCCAACACTTGTCACTACTTTCTCTTATGGTGTTCAATGCTTTTCAAGATACCCAGATCATATGAAGCGGCACGACTTCTTCAAGAGCGCCATGCCTGAGGGATACGTGCAGGAGAGGACCATCTTCTTCAAGGACGACGGGAACTACAAGACACGTGCTGAAGTCAAGTTTGAGGGAGACACCCTCGTCAACAGGATCGAGCTTAAGGGAATCGATTTCAAGGAGGACGGAAACATCCTCGGCCACAAGTTGGAATACAACTACAACTCCCACAACGTATACATCATGGCCGACAAGCAAAAGAACGGCATCAAAGCCAACTTCAAGACCCGCCACAACATCGAAGACGGCGGCGTGCAACTCGCTGATCATTATCAACAAAATACTCCAATTGGCGATGGCCCTGTCCTTTTACCAGACAACCATTACCTGTCCACACAATCTGCCCTTTCGAAAGATCCCAACGAAAAGAGAGACCACATGGTCCTTCTTGAGTTTGTAACAGCTGCTGGGATTACACATGGCATGGATGAACTATACAAACATGATGAGCTTTAAGACGTCCGATCGTTCAAACATTTGGCAATAAAGTTTCTTAAGATTGAATCCTGTTGCCGGTCTTGCGATGATTATCATATAATTTCTGTTGAATTACGTTAAGCATGTAATAATTAACATGTAATGCATGACGTTATTTATGAGATGGGTTTTTATGATTAGAGTCCCGCAATTATACATTTAATACGCGATAGAAAACAAAATATAGCGCGCAAACTAGGATAAATTATCGCGCGCGGTGTCATCTATGTTACTAGATCGGGAATTGATCCCCCCTCGACAGCTTCCGGAAAGGGCGAATTCGCAACTTTGTATACAAAAGTTGCCGAGCTCGAATTCGTAATCATGGTCATAGCTGTTTCCTGTGTGAAATTGTTATCCGCTCACAATTCCACACAACATACGAGCCGGAAGCATAAAGTGTAAAGCCTGGGGTGCCTAATGAGTGAGCTAACTCACATTAATTGCGTTGCGCTCACTGCCCGCTTTCCAGTCGGGAAACCTGTCGTGCCAGCTGCATTAATGAATCGGCCAACGCGCGGGGAGAGGCGGTTTGCGTATTGGCTAGAGCAGCTTGCCAACATGGTGGAGCACGACACTCTCGTCTACTCCAAGAATATCAAAGATACAGTCTCAGAAGACCAAAGGGCTATTGAGACTTTTCAACAAAGGGTAATATCGGGAAACCTCCTCGGATTCCATTGCCCAGCTATCTGTCACTTCATCAAAAGGACAGTAGAAAAGGAAGGTGGCACCTACAAATGCCATCATTGCGATAAAGGAAAGGCTATCGTTCAAGATGCCTCTGCCGACAGTGGTCCCAAAGATGGACCCCCACCCACGAGGAGCATCGTGGAAAAAGAAGACGTTCCAACCACGTCTTCAAAGCAAGTGGATTGATGTGATAACATGGTGGAGCACGACACTCTCGTCTACTCCAAGAATATCAAAGATACAGTCTCAGAAGACCAAAGGGCTATTGAGACTTTTCAACAAAGGGTAATATCGGGAAACCTCCTCGGATTCCATTGCCCAGCTATCTGTCACTTCATCAAAAGGACAGTAGAAAAGGAAGGTGGCACCTACAAATGCCATCATTGCGATAAAGGAAAGGCTATCGTTCAAGATGCCTCTGCCGACAGTGGTCCCAAAGATGGACCCCCACCCACGAGGAGCATCGTGGAAAAAGAAGACGTTCCAACCACGTCTTCAAAGCAAGTGGATTGATGTGATATCTCCACTGACGTAAGGGATGACGCACAATCCCACTATCCTTCGCAAGACCTTCCTCTATATAAGGAAGTTCATTTCATTTGGAGAGGACACGCTGAAATCACCAGTCTCTCTCTACAAATCTATCTCTCTCGAGCTTTCGCAGATCCGGGGGGGCAATGAGATATGAAAAAGCCTGAACTCACCGCGACGTCTGTCGAGAAGTTTCTGATCGAAAAGTTCGACAGCGTCTCCGACCTGATGCAGCTCTCGGAGGGCGAAGAATCTCGTGCTTTCAGCTTCGATGTAGGAGGGCGTGGATATGTCCTGCGGGTAAATAGCTGCGCCGATGGTTTCTACAAAGATCGTTATGTTTATCGGCACTTTGCATCGGCCGCGCTCCCGATTCCGGAAGTGCTTGACATTGGGGAGTTTAGCGAGAGCCTGACCTATTGCATCTCCCGCCGTGCACAGGGTGTCACGTTGCAAGACCTGCCTGAAACCGAACTGCCCGCTGTTCTACAACCGGTCGCGGAGGCTATGGATGCGATCGCTGCGGCCGATCTTAGCCAGACGAGCGGGTTCGGCCCATTCGGACCGCAAGGAATCGGTCAATACACTACATGGCGTGATTTCATATGCGCGATTGCTGATCCCCATGTGTATCACTGGCAAACTGTGATGGACGACACCGTCAGTGCGTCCGTCGCGCAGGCTCTCGATGAGCTGATGCTTTGGGCCGAGGACTGCCCCGAAGTCCGGCACCTCGTGCACGCGGATTTCGGCTCCAACAATGTCCTGACGGACAATGGCCGCATAACAGCGGTCATTGACTGGAGCGAGGCGATGTTCGGGGATTCCCAATACGAGGTCGCCAACATCTTCTTCTGGAGGCCGTGGTTGGCTTGTATGGAGCAGCAGACGCGCTACTTCGAGCGGAGGCATCCGGAGCTTGCAGGATCGCCACGACTCCGGGCGTATATGCTCCGCATTGGTCTTGACCAACTCTATCAGAGCTTGGTTGACGGCAATTTCGATGATGCAGCTTGGGCGCAGGGTCGATGCGACGCAATCGTCCGATCCGGAGCCGGGACTGTCGGGCGTACACAAATCGCCCGCAGAAGCGCGGCCGTCTGGACCGATGGCTGTGTAGAAGTACTCGCCGATAGTGGAAACCGACGCCCCAGCACTCGTCCGAGGGCAAAGAAATAGAGTAGATGCCGACGCGGATCTGTCGATCGACAAGCTCGAGTTTCTCCATAATAATGTGTGAGTAGTTCCCAGATAAGGGAATTAGGGTTCCTATAGGGTTTCGCTCATGTGTTGAGCATATAAGAAACCCTTAGTATGTATTTGTATTTGTAAAATACTTCTATCAATAAAATTTCTAATTCCTAAAACCAAAATCCAGTACTAAAATCCAGATCCCCCGAATTAATTCGGCGTTAATTCAGTACATTAAAAACGTCCGCAATGTGTTATTAAGTTGTCTAAGCGTCAATTTGTTTACACCACAATATATCCTGCCACCAGCCAGCCAACAGCTCCCCGACCGGCAGCTCGGCACAAAATCACCACTCGATACAGGCAGCCCATCAGTCCGGGACGGCGTCAGCGGGAGAGCCGTTGTAAGGCGGCAGACTTTGCTCATGTTACCGATGCTATTCGGAAGAACGGCAACTAAGCTGCCGGGTTTGAAACACGGATGATCTCGCGGAGGGTAGCATGTTGATTGTAACGATGACAGAGCGTTGCTGCCTGTGATCACCGCGGTTTCAAAATCGGCTCCGTCGATACTATGTTATACGCCAACTTTGAAAACAACTTTGAAAAAGCTGTTTTCTGGTATTTAAGGTTTTAGAATGCAAGGAACAGTGAATTGGAGTTCGTCTTGTTATAATTAGCTTCTTGGGGTATCTTTAAATACTGTAGAAAAGAGGAAGGAAATAATAAATGGCTAAAATGAGAATATCACCGGAATTGAAAAAACTGATCGAAAAATACCGCTGCGTAAAAGATACGGAAGGAATGTCTCCTGCTAAGGTATATAAGCTGGTGGGAGAAAATGAAAACCTATATTTAAAAATGACGGACAGCCGGTATAAAGGGACCACCTATGATGTGGAACGGGAAAAGGACATGATGCTATGGCTGGAAGGAAAGCTGCCTGTTCCAAAGGTCCTGCACTTTGAACGGCATGATGGCTGGAGCAATCTGCTCATGAGTGAGGCCGATGGCGTCCTTTGCTCGGAAGAGTATGAAGATGAACAAAGCCCTGAAAAGATTATCGAGCTGTATGCGGAGTGCATCAGGCTCTTTCACTCCATCGACATATCGGATTGTCCCTATACGAATAGCTTAGACAGCCGCTTAGCCGAATTGGATTACTTACTGAATAACGATCTGGCCGATGTGGATTGCGAAAACTGGGAAGAAGACACTCCATTTAAAGATCCGCGCGAGCTGTATGATTTTTTAAAGACGGAAAAGCCCGAAGAGGAACTTGTCTTTTCCCACGGCGACCTGGGAGACAGCAACATCTTTGTGAAAGATGGCAAAGTAAGTGGCTTTATTGATCTTGGGAGAAGCGGCAGGGCGGACAAGTGGTATGACATTGCCTTCTGCGTCCGGTCGATCAGGGAGGATATCGGGGAAGAACAGTATGTCGAGCTATTTTTTGACTTACTGGGGATCAAGCCTGATTGGGAGAAAATAAAATATTATATTTTACTGGATGAATTGTTTTAGTACCTAGAATGCATGACCAAAATCCCTTAACGTGAGTTTTCGTTCCACTGAGCGTCAGACCCCGTAGAAAAGATCAAAGGATCTTCTTGAGATCCTTTTTTTCTGCGCGTAATCTGCTGCTTGCAAACAAAAAAACCACCGCTACCAGCGGTGGTTTGTTTGCCGGATCAAGAGCTACCAACTCTTTTTCCGAAGGTAACTGGCTTCAGCAGAGCGCAGATACCAAATACTGTCCTTCTAGTGTAGCCGTAGTTAGGCCACCACTTCAAGAACTCTGTAGCACCGCCTACATACCTCGCTCTGCTAATCCTGTTACCAGTGGCTGCTGCCAGTGGCGATAAGTCGTGTCTTACCGGGTTGGACTCAAGACGATAGTTACCGGATAAGGCGCAGCGGTCGGGCTGAACGGGGGGTTCGTGCACACAGCCCAGCTTGGAGCGAACGACCTACACCGAACTGAGATACCTACAGCGTGAGCTATGAGAAAGCGCCACGCTTCCCGAAGGGAGAAAGGCGGACAGGTATCCGGTAAGCGGCAGGGTCGGAACAGGAGAGCGCACGAGGGAGCTTCCAGGGGGAAACGCCTGGTATCTTTATAGTCCTGTCGGGTTTCGCCACCTCTGACTTGAGCGTCGATTTTTGTGATGCTCGTCAGGGGGGCGGAGCCTATGGAAAAACGCCAGCAACGCGGCCTTTTTACGGTTCCTGGCCTTTTGCTGGCCTTTTGCTCACATGTTCTTTCCTGCGTTATCCCCTGATTCTGTGGATAACCGTATTACCGCCTTTGAGTGAGCTGATACCGCTCGCCGCAGCCGAACGACCGAGCGCAGCGAGTCAGTGAGCGAGGAAGCGGAAGAGCGCCTGATGCGGTATTTTCTCCTTACGCATCTGTGCGGTATTTCACACCGCATATGGTGCACTCTCAGTACAATCTGCTCTGATGCCGCATAGTTAAGCCAGTATACACTCCGCTATCGCTACGTGACTGGGTCATGGCTGCGCCCCGACACCCGCCAACACCCGCTGACGCGCCCTGACGGGCTTGTCTGCTCCCGGCATCCGCTTACAGACAAGCTGTGACCGTCTCCGGGAGCTGCATGTGTCAGAGGTTTTCACCGTCATCACCGAAACGCGCGAGGCAGGGTGCCTTGATGTGGGCGCCGGCGGTCGAGTGGCGACGGCGCGGCTTGTCCGCGCCCTGGTAGATTGCCTGGCCGTAGGCCAGCCATTTTTGAGCGGCCAGCGGCCGCGATAGGCCGACGCGAAGCGGCGGGGCGTAGGGAGCGCAGCGACCGAAGGGTAGGCGCTTTTTGCAGCTCTTCGGCTGTGCGCTGGCCAGACAGTTATGCACAGGCCAGGCGGGTTTTAAGAGTTTTAATAAGTTTTAAAGAGTTTTAGGCGGAAAAATCGCCTTTTTTCTCTTTTATATCAGTCACTTACATGTGTGACCGGTTCCCAATGTACGGCTTTGGGTTCCCAATGTACGGGTTCCGGTTCCCAATGTACGGCTTTGGGTTCCCAATGTACGTGCTATCCACAGGAAAGAGACCTTTTCGACCTTTTTCCCCTGCTAGGGCAATTTGCCCTAGCATCTGCTCCGTACATTAGGAACCGGCGGATGCTTCGCCCTCGATCAGGTTGCGGTAGCGCATGACTAGGATCGGGCCAGCCTGCCCCGCCTCCTCCTTCAAATCGTACTCCGGCAGGTCATTTGACCCGATCAGCTTGCGCACGGTGAAACAGAACTTCTTGAACTCTCCGGCGCTGCCACTGCGTTCGTAGATCGTCTTGAACAACCATCTGGCTTCTGCCTTGCCTGCGGCGCGGCGTGCCAGGCGGTAGAGAAAACGGCCGATGCCGGGATCGATCAAAAAGTAATCGGGGTGAACCGTCAGCACGTCCGGGTTCTTGCCTTCTGTGATCTCGCGGTACATCCAATCAGCTAGCTCGATCTCGATGTACTCCGGCCGCCCGGTTTCGCTCTTTACGATCTTGTAGCGGCTAATCAAGGCTTCACCCTCGGATACCGTCACCAGGCGGCCGTTCTTGGCCTTCTTCGTACGCTGCATGGCAACGTGCGTGGTGTTTAACCGAATGCAGGTTTCTACCAGGTCGTCTTTCTGCTTTCCGCCATCGGCTCGCCGGCAGAACTTGAGTACGTCCGCAACGTGTGGACGGAACACGCGGCCGGGCTTGTCTCCCTTCCCTTCCCGGTATCGGTTCATGGATTCGGTTAGATGGGAAACCGCCATCAGTACCAGGTCGTAATCCCACACACTGGCCATGCCGGCCGGCCCTGCGGAAACCTCTACGTGCCCGTCTGGAAGCTCGTAGCGGATCACCTCGCCAGCTCGTCGGTCACGCTTCGACAGACGGAAAACGGCCACGTCCATGATGCTGCGACTATCGCGGGTGCCCACGTCATAGAGCATCGGAACGAAAAAATCTGGTTGCTCGTCGCCCTTGGGCGGCTTCCTAATCGACGGCGCACCGGCTGCCGGCGGTTGCCGGGATTCTTTGCGGATTCGATCAGCGGCCGCTTGCCACGATTCACCGGGGCGTGCTTCTGCCTCGATGCGTTGCCGCTGGGCGGCCTGCGCGGCCTTCAACTTCTCCACCAGGTCATCACCCAGCGCCGCGCCGATTTGTACCGGGCCGGATGGTTTGCGACCGTCACGCCGATTCCTCGGGCTTGGGGGTTCCAGTGCCATTGCAGGGCCGGCAGACAACCCAGCCGCTTACGCCTGGCCAACCGCCCGTTCCTCCACACATGGGGCATTCCACGGCGTCGGTGCCTGGTTGTTCTTGATTTTCCATGCCGCCTCCTTTAGCCGCTAAAATTCATCTACTCATTTATTCATTTGCTCATTTACTCTGGTAGCTGCGCGATGTATTCAGATAGCAGCTCGGTAATGGTCTTGCCTTGGCGTACCGCGTACATCTTCAGCTTGGTGTGATCCTCCGCCGGCAACTGAAAGTTGACCCGCTTCATGGCTGGCGTGTCTGCCAGGCTGGCCAACGTTGCAGCCTTGCTGCTGCGTGCGCTCGGACGGCCGGCACTTAGCGTGTTTGTGCTTTTGCTCATTTTCTCTTTACCTCATTAACTCAAATGAGTTTTGATTTAATTTCAGCGGCCAGCGCCTGGACCTCGCGGGCAGCGTCGCCCTCGGGTTCTGATTCAAGAACGGTTGTGCCGGCGGCGGCAGTGCCTGGGTAGCTCACGCGCTGCGTGATACGGGACTCAAGAATGGGCAGCTCGTACCCGGCCAGCGCCTCGGCAACCTCACCGCCGATGCGCGTGCCTTTGATCGCCCGCGACACGACAAAGGCCGCTTGTAGCCTTCCATCCGTGACCTCAATGCGCTGCTTAACCAGCTCCACCAGGTCGGCGGTGGCCCATATGTCGTAAGGGCTTGGCTGCACCGGAATCAGCACGAAGTCGGCTGCCTTGATCGCGGACACAGCCAAGTCCGCCGCCTGGGGCGCTCCGTCGATCACTACGAAGTCGCGCCGGCCGATGGCCTTCACGTCGCGGTCAATCGTCGGGCGGTCGATGCCGACAACGGTTAGCGGTTGATCTTCCCGCACGGCCGCCCAATCGCGGGCACTGCCCTGGGGATCGGAATCGACTAACAGAACATCGGCCCCGGCGAGTTGCAGGGCGCGGGCTAGATGGGTTGCGATGGTCGTCTTGCCTGACCCGCCTTTCTGGTTAAGTACAGCGATAACCTTCATGCGTTCCCCTTGCGTATTTGTTTATTTACTCATCGCATCATATACGCAGCGACCGCATGACGCAAGCTGTTTTACTCAAATACACATCACCTTTTTAGACGGCGGCGCTCGGTTTCTTCAGCGGCCAAGCTGGCCGGCCAGGCCGCCAGCTTGGCATCAGACAAACCGGCCAGGATTTCATGCAGCCGCACGGTTGAGACGTGCGCGGGCGGCTCGAACACGTACCCGGCCGCGATCATCTCCGCCTCGATCTCTTCGGTAATGAAAAACGGTTCGTCCTGGCCGTCCTGGTGCGGTTTCATGCTTGTTCCTCTTGGCGTTCATTCTCGGCGGCCGCCAGGGCGTCGGCCTCGGTCAATGCGTCCTCACGGAAGGCACCGCGCCGCCTGGCCTCGGTGGGCGTCACTTCCTCGCTGCGCTCAAGTGCGCGGTACAGGGTCGAGCGATGCACGCCAAGCAGTGCAGCCGCCTCTTTCACGGTGCGGCCTTCCTGGTCGATCAGCTCGCGGGCGTGCGCGATCTGTGCCGGGGTGAGGGTAGGGCGGGGGCCAAACTTCACGCCTCGGGCCTTGGCGGCCTCGCGCCCGCTCCGGGTGCGGTCGATGATTAGGGAACGCTCGAACTCGGCAATGCCGGCGAACACGGTCAACACCATGCGGCCGGCCGGCGTGGTGGTGTCGGCCCACGGCTCTGCCAGGCTACGCAGGCCCGCGCCGGCCTCCTGGATGCGCTCGGCAATGTCCAGTAGGTCGCGGGTGCTGCGGGCCAGGCGGTCTAGCCTGGTCACTGTCACAACGTCGCCAGGGCGTAGGTGGTCAAGCATCCTGGCCAGCTCCGGGCGGTCGCGCCTGGTGCCGGTGATCTTCTCGGAAAACAGCTTGGTGCAGCCGGCCGCGTGCAGTTCGGCCCGTTGGTTGGTCAAGTCCTGGTCGTCGGTGCTGACGCGGGCATAGCCCAGCAGGCCAGCGGCGGCGCTCTTGTTCATGGCGTAATGTCTCCGGTTCTAGTCGCAAGTATTCTACTTTATGCGACTAAAACACGCGACAAGAAAACGCCAGGAAAAGGGCAGGGCGGCAGCCTGTCGCGTAACTTAGGACTTGTGCGACATGTCGTTTTCAGAAGACGGCTGCACTGAACGTCAGAAGCCGACTGCACTATAGCAGCGGAGGGGTTGGATCAAAGTACTTTGATCCCGAGGGGAACCCTGTGGTTGGCATGCACATACAAATGGACGAACGGATAAACCTTTTCACGCCCTTTTAAATATCCGTTATTCTAATAAACGCTCTTTTCTCTTAGGTTTACCCGCCAATATATCCTGTCAAACACTGATAGTTTAAACTGAAGGCGGGAAACGACAATCTGATCCAAGCTCAAGCTGCTCTAGCATTCGCCATTCAGGCTGCGCAACTGTTGGGAAGGGCGATCGGTGCGGGCCTCTTCGCTATTACGCCAGCTGGCGAAAGGGGGATGTGCTGCAAGGCGATTAAGTTGGGTAACGCCAGGGTTTTCCCAGTCACGACGTTGTAAAACGACGGCCAGTGCCAAGCTTTGAGACTTTTCAACAAAGGGTAATATCGGGAAACCTCCTCGGATTCCATTGCCCAGCTATCTGTCACTTCATCAAAAGGACAGTAGAAAAGGAAGGTGGCACCTACAAATGCCATCATTGCGATAAAGGAAAGGCTATCGTTCAAGATGCCTCTGCCGACAGTGGTCCCAAAGATGGACCCCCACCCACGAGGAGCATCGTGGAAAAAGAAGACGTTCCAACCACGTCTTCAAAGCAAGTGGATTGATGTGATAACATGGTGGAGCACGACACTCTCGTCTACTCCAAGAATATCAAAGATACAGTCTCAGAAGACCAAAGGGCTATTGAGACTTTTCAACAAAGGGTAATATCGGGAAACCTCCTCGGATTCCATTGCCCAGCTATCTGTCACTTCATCAAAAGGACAGTAGAAAAGGAAGGTGGCACCTACAAATGCCATCATTGCGATAAAGGAAAGGCTATCGTTCAAGATGCCTCTGCCGACAGTGGTCCCAAAGATGGACCCCCACCCACGAGGAGCATCGTGGAAAAAGAAGACGTTCCAACCACGTCTTCAAAGCAAGTGGATTGATGTGATATCTCCACTGACGTAAGGGATGACGCACAATCCCACTATCCTTCGCAAGACCTTCCTCTATATAAGGAAGTTCATTTCATTTGGAGAGGACACGCTGACTGCA

**Sequences S5:** DNA sequence of the vector pTarget: **p35S-pT7-ALS** (p35S_pT7_ALS_T7term_polA) (two-vector system)

GTACAACAGCGACTACAAGGATGACGATGACAAGGCTTAGAGCTCGAATTTCCCCGATCGTTCAAACATTTGGCAATAAAGTTTCTTAAGATTGAATCCTGTTGCCGGTCTTGCGATGATTATCATATAATTTCTGTTGAATTACGTTAAGCATGTAATAATTAACATGTAATGCATGACGTTATTTATGAGATGGGTTTTTATGATTAGAGTCCCGCAATTATACATTTAATACGCGATAGAAAACAAAATATAGCGCGCAAACTAGGATAAATTATCGCGCGCGGTGTCATCTATGTTACTAGATCGGGAATTCACTGGCCGTCGTTTTACACTGGCCGTCGTTTTACAACGTCGTGACTGGGAAAACCCTGGCGTTACCCAACTTAATCGCCTTGCAGCACATCCCCCTTTCGCCAGCTGGCGTAATAGCGAAGAGGCCCGCACCGATCGCCCTTCCCAACAGTTGCGCAGCCTGAATGGCGAATGCTAGAGCAGCTTGAGCTTGGATCAGATTGTCGTTTCCCGCCTTCAGTTTAAACTATCAGTGTTTGACAGGATATATTGGCGGGTAAACCTAAGAGAAAAGAGCGTTTATTAGAATAACGGATATTTAAAAGGGCGTGAAAAGGTTTATCCGTTCGTCCATTTGTATGTGCATGCCAACCACAGGGTTCCCCTCGGGATCAAAGTACTTTGATCCAACCCCTCCGCTGCTATAGTGCAGTCGGCTTCTGACGTTCAGTGCAGCCGTCTTCTGAAAACGACATGTCGCACAAGTCCTAAGTTACGCGACAGGCTGCCGCCCTGCCCTTTTCCTGGCGTTTTCTTGTCGCGTGTTTTAGTCGCATAAAGTAGAATACTTGCGACTAGAACCGGAGACATTACGCCATGAACAAGAGCGCCGCCGCTGGCCTGCTGGGCTATGCCCGCGTCAGCACCGACGACCAGGACTTGACCAACCAACGGGCCGAACTGCACGCGGCCGGCTGCACCAAGCTGTTTTCCGAGAAGATCACCGGCACCAGGCGCGACCGCCCGGAGCTGGCCAGGATGCTTGACCACCTACGCCCTGGCGACGTTGTGACAGTGACCAGGCTAGACCGCCTGGCCCGCAGCACCCGCGACCTACTGGACATTGCCGAGCGCATCCAGGAGGCCGGCGCGGGCCTGCGTAGCCTGGCAGAGCCGTGGGCCGACACCACCACGCCGGCCGGCCGCATGGTGTTGACCGTGTTCGCCGGCATTGCCGAGTTCGAGCGTTCCCTAATCATCGACCGCACCCGGAGCGGGCGCGAGGCCGCCAAGGCCCGAGGCGTGAAGTTTGGCCCCCGCCCTACCCTCACCCCGGCACAGATCGCGCACGCCCGCGAGCTGATCGACCAGGAAGGCCGCACCGTGAAAGAGGCGGCTGCACTGCTTGGCGTGCATCGCTCGACCCTGTACCGCGCACTTGAGCGCAGCGAGGAAGTGACGCCCACCGAGGCCAGGCGGCGCGGTGCCTTCCGTGAGGACGCATTGACCGAGGCCGACGCCCTGGCGGCCGCCGAGAATGAACGCCAAGAGGAACAAGCATGAAACCGCACCAGGACGGCCAGGACGAACCGTTTTTCATTACCGAAGAGATCGAGGCGGAGATGATCGCGGCCGGGTACGTGTTCGAGCCGCCCGCGCACGTCTCAACCGTGCGGCTGCATGAAATCCTGGCCGGTTTGTCTGATGCCAAGCTGGCGGCCTGGCCGGCCAGCTTGGCCGCTGAAGAAACCGAGCGCCGCCGTCTAAAAAGGTGATGTGTATTTGAGTAAAACAGCTTGCGTCATGCGGTCGCTGCGTATATGATGCGATGAGTAAATAAACAAATACGCAAGGGGAACGCATGAAGGTTATCGCTGTACTTAACCAGAAAGGCGGGTCAGGCAAGACGACCATCGCAACCCATCTAGCCCGCGCCCTGCAACTCGCCGGGGCCGATGTTCTGTTAGTCGATTCCGATCCCCAGGGCAGTGCCCGCGATTGGGCGGCCGTGCGGGAAGATCAACCGCTAACCGTTGTCGGCATCGACCGCCCGACGATTGACCGCGACGTGAAGGCCATCGGCCGGCGCGACTTCGTAGTGATCGACGGAGCGCCCCAGGCGGCGGACTTGGCTGTGTCCGCGATCAAGGCAGCCGACTTCGTGCTGATTCCGGTGCAGCCAAGCCCTTACGACATATGGGCCACCGCCGACCTGGTGGAGCTGGTTAAGCAGCGCATTGAGGTCACGGATGGAAGGCTACAAGCGGCCTTTGTCGTGTCGCGGGCGATCAAAGGCACGCGCATCGGCGGTGAGGTTGCCGAGGCGCTGGCCGGGTACGAGCTGCCCATTCTTGAGTCCCGTATCACGCAGCGCGTGAGCTACCCAGGCACTGCCGCCGCCGGCACAACCGTTCTTGAATCAGAACCCGAGGGCGACGCTGCCCGCGAGGTCCAGGCGCTGGCCGCTGAAATTAAATCAAAACTCATTTGAGTTAATGAGGTAAAGAGAAAATGAGCAAAAGCACAAACACGCTAAGTGCCGGCCGTCCGAGCGCACGCAGCAGCAAGGCTGCAACGTTGGCCAGCCTGGCAGACACGCCAGCCATGAAGCGGGTCAACTTTCAGTTGCCGGCGGAGGATCACACCAAGCTGAAGATGTACGCGGTACGCCAAGGCAAGACCATTACCGAGCTGCTATCTGAATACATCGCGCAGCTACCAGAGTAAATGAGCAAATGAATAAATGAGTAGATGAATTTTAGCGGCTAAAGGAGGCGGCATGGAAAATCAAGAACAACCAGGCACCGACGCCGTGGAATGCCCCATGTGTGGAGGAACGGGCGGTTGGCCAGGCGTAAGCGGCTGGGTTGTCTGCCGGCCCTGCAATGGCACTGGAACCCCCAAGCCCGAGGAATCGGCGTGACGGTCGCAAACCATCCGGCCCGGTACAAATCGGCGCGGCGCTGGGTGATGACCTGGTGGAGAAGTTGAAGGCCGCGCAGGCCGCCCAGCGGCAACGCATCGAGGCAGAAGCACGCCCCGGTGAATCGTGGCAAGCGGCCGCTGATCGAATCCGCAAAGAATCCCGGCAACCGCCGGCAGCCGGTGCGCCGTCGATTAGGAAGCCGCCCAAGGGCGACGAGCAACCAGATTTTTTCGTTCCGATGCTCTATGACGTGGGCACCCGCGATAGTCGCAGCATCATGGACGTGGCCGTTTTCCGTCTGTCGAAGCGTGACCGACGAGCTGGCGAGGTGATCCGCTACGAGCTTCCAGACGGGCACGTAGAGGTTTCCGCAGGGCCGGCCGGCATGGCCAGTGTGTGGGATTACGACCTGGTACTGATGGCGGTTTCCCATCTAACCGAATCCATGAACCGATACCGGGAAGGGAAGGGAGACAAGCCCGGCCGCGTGTTCCGTCCACACGTTGCGGACGTACTCAAGTTCTGCCGGCGAGCCGATGGCGGAAAGCAGAAAGACGACCTGGTAGAAACCTGCATTCGGTTAAACACCACGCACGTTGCCATGCAGCGTACGAAGAAGGCCAAGAACGGCCGCCTGGTGACGGTATCCGAGGGTGAAGCCTTGATTAGCCGCTACAAGATCGTAAAGAGCGAAACCGGGCGGCCGGAGTACATCGAGATCGAGCTAGCTGATTGGATGTACCGCGAGATCACAGAAGGCAAGAACCCGGACGTGCTGACGGTTCACCCCGATTACTTTTTGATCGATCCCGGCATCGGCCGTTTTCTCTACCGCCTGGCACGCCGCGCCGCAGGCAAGGCAGAAGCCAGATGGTTGTTCAAGACGATCTACGAACGCAGTGGCAGCGCCGGAGAGTTCAAGAAGTTCTGTTTCACCGTGCGCAAGCTGATCGGGTCAAATGACCTGCCGGAGTACGATTTGAAGGAGGAGGCGGGGCAGGCTGGCCCGATCCTAGTCATGCGCTACCGCAACCTGATCGAGGGCGAAGCATCCGCCGGTTCCTAATGTACGGAGCAGATGCTAGGGCAAATTGCCCTAGCAGGGGAAAAAGGTCGAAAAGCACTCTTTCCTGTGGATAGCACGTACATTGGGAACCCAAAGCCGTACATTGGGAACCGGAACCCGTACATTGGGAACCCAAAGCCGTACATTGGGAACCGGTCACACATGTAAGTGACTGATATAAAAGAGAAAAAAGGCGATTTTTCCGCCTAAAACTCTTTAAAACTTATTAAAACTCTTAAAACCCGCCTGGCCTGTGCATAACTGTCTGGCCAGCGCACAGCCGAAGAGCTGCAAAAAGCGCCTACCCTTCGGTCGCTGCGCTCCCTACGCCCCGCCGCTTCGCGTCGGCCTATCGCGGCCGCTGGCCGCTCAAAAATGGCTGGCCTACGGCCAGGCAATCTACCAGGGCGCGGACAAGCCGCGCCGTCGCCACTCGACCGCCGGCGCCCACATCAAGGCACCCTGCCTCGCGCGTTTCGGTGATGACGGTGAAAACCTCTGACACATGCAGCTCCCGGAGACGGTCACAGCTTGTCTGTAAGCGGATGCCGGGAGCAGACAAGCCCGTCAGGGCGCGTCAGCGGGTGTTGGCGGGTGTCGGGGCGCAGCCATGACCCAGTCACGTAGCGATAGCGGAGTGTATACTGGCTTAACTATGCGGCATCAGAGCAGATTGTACTGAGAGTGCACCATATGCGGTGTGAAATACCGCACAGATGCGTAAGGAGAAAATACCGCATCAGGCGCTCTTCCGCTTCCTCGCTCACTGACTCGCTGCGCTCGGTCGTTCGGCTGCGGCGAGCGGTATCAGCTCACTCAAAGGCGGTAATACGGTTATCCACAGAATCAGGGGATAACGCAGGAAAGAACATGTGAGCAAAAGGCCAGCAAAAGGCCAGGAACCGTAAAAAGGCCGCGTTGCTGGCGTTTTTCCATAGGCTCCGCCCCCCTGACGAGCATCACAAAAATCGACGCTCAAGTCAGAGGTGGCGAAACCCGACAGGACTATAAAGATACCAGGCGTTTCCCCCTGGAAGCTCCCTCGTGCGCTCTCCTGTTCCGACCCTGCCGCTTACCGGATACCTGTCCGCCTTTCTCCCTTCGGGAAGCGTGGCGCTTTCTCATAGCTCACGCTGTAGGTATCTCAGTTCGGTGTAGGTCGTTCGCTCCAAGCTGGGCTGTGTGCACGAACCCCCCGTTCAGCCCGACCGCTGCGCCTTATCCGGTAACTATCGTCTTGAGTCCAACCCGGTAAGACACGACTTATCGCCACTGGCAGCAGCCACTGGTAACAGGATTAGCAGAGCGAGGTATGTAGGCGGTGCTACAGAGTTCTTGAAGTGGTGGCCTAACTACGGCTACACTAGAAGGACAGTATTTGGTATCTGCGCTCTGCTGAAGCCAGTTACCTTCGGAAAAAGAGTTGGTAGCTCTTGATCCGGCAAACAAACCACCGCTGGTAGCGGTGGTTTTTTTGTTTGCAAGCAGCAGATTACGCGCAGAAAAAAAGGATCTCAAGAAGATCCTTTGATCTTTTCTACGGGGTCTGACGCTCAGTGGAACGAAAACTCACGTTAAGGGATTTTGGTCATGCATTCTAGGTACTAAAACAATTCATCCAGTAAAATATAATATTTTATTTTCTCCCAATCAGGCTTGATCCCCAGTAAGTCAAAAAATAGCTCGACATACTGTTCTTCCCCGATATCCTCCCTGATCGACCGGACGCAGAAGGCAATGTCATACCACTTGTCCGCCCTGCCGCTTCTCCCAAGATCAATAAAGCCACTTACTTTGCCATCTTTCACAAAGATGTTGCTGTCTCCCAGGTCGCCGTGGGAAAAGACAAGTTCCTCTTCGGGCTTTTCCGTCTTTAAAAAATCATACAGCTCGCGCGGATCTTTAAATGGAGTGTCTTCTTCCCAGTTTTCGCAATCCACATCGGCCAGATCGTTATTCAGTAAGTAATCCAATTCGGCTAAGCGGCTGTCTAAGCTATTCGTATAGGGACAATCCGATATGTCGATGGAGTGAAAGAGCCTGATGCACTCCGCATACAGCTCGATAATCTTTTCAGGGCTTTGTTCATCTTCATACTCTTCCGAGCAAAGGACGCCATCGGCCTCACTCATGAGCAGATTGCTCCAGCCATCATGCCGTTCAAAGTGCAGGACCTTTGGAACAGGCAGCTTTCCTTCCAGCCATAGCATCATGTCCTTTTCCCGTTCCACATCATAGGTGGTCCCTTTATACCGGCTGTCCGTCATTTTTAAATATAGGTTTTCATTTTCTCCCACCAGCTTATATACCTTAGCAGGAGACATTCCTTCCGTATCTTTTACGCAGCGGTATTTTTCGATCAGTTTTTTCAATTCCGGTGATATTCTCATTTTAGCCATTTATTATTTCCTTCCTCTTTTCTACAGTATTTAAAGATACCCCAAGAAGCTAATTATAACAAGACGAACTCCAATTCACTGTTCCTTGCATTCTAAAACCTTAAATACCAGAAAACAGCTTTTTCAAAGTTGTTTTCAAAGTTGGCGTATAACATAGTATCGACGGAGCCGATTTTGAAACCGCGGTGATCACAGGCAGCAACGCTCTGTCATCGTTACAATCAACATGCTACCCTCCGCGAGATCATCCGTGTTTCAAACCCGGCAGCTTAGTTGCCGTTCTTCCGAATAGCATCGGTAACATGAGCAAAGTCTGCCGCCTTACAACGGCTCTCCCGCTGACGCCGTCCCGGACTGATGGGCTGCCTGTATCGAGTGGTGATTTTGTGCCGAGCTGCCGGTCGGGGAGCTGTTGGCTGGCTGGTGGCAGGATATATTGTGGTGTAAACAAATTGACGCTTAGACAACTTAATAACACATTGCGGACGTTTTTAATGTACTGAATTAACGCCGAATTAATTCGGGGGATCTGGATTTTAGTACTGGATTTTGGTTTTAGGAATTAGAAATTTTATTGATAGAAGTATTTTACAAATACAAATACATACTAAGGGTTTCTTATATGCTCAACACATGAGCGAAACCCTATAGGAACCCTAATTCCCTTATCTGGGAACTACTCACACATTATTATGGAGAAACTCGAGCTTGTCGATCGACAGATCCGGTCGGCATCTACTCTATTTCTTTGCCCTCGGACGAGTGCTGGGGCGTCGGTTTCCACTATCGGCGAGTACTTCTACACAGCCATCGGTCCAGACGGCCGCGCTTCTGCGGGCGATTTGTGTACGCCCGACAGTCCCGGCTCCGGATCGGACGATTGCGTCGCATCGACCCTGCGCCCAAGCTGCATCATCGAAATTGCCGTCAACCAAGCTCTGATAGAGTTGGTCAAGACCAATGCGGAGCATATACGCCCGGAGTCGTGGCGATCCTGCAAGCTCCGGATGCCTCCGCTCGAAGTAGCGCGTCTGCTGCTCCATACAAGCCAACCACGGCCTCCAGAAGAAGATGTTGGCGACCTCGTATTGGGAATCCCCGAACATCGCCTCGCTCCAGTCAATGACCGCTGTTATGCGGCCATTGTCCGTCAGGACATTGTTGGAGCCGAAATCCGCGTGCACGAGGTGCCGGACTTCGGGGCAGTCCTCGGCCCAAAGCATCAGCTCATCGAGAGCCTGCGCGACGGACGCACTGACGGTGTCGTCCATCACAGTTTGCCAGTGATACACATGGGGATCAGCAATCGCGCATATGAAATCACGCCATGTAGTGTATTGACCGATTCCTTGCGGTCCGAATGGGCCGAACCCGCTCGTCTGGCTAAGATCGGCCGCAGCGATCGCATCCATAGCCTCCGCGACCGGTTGTAGAACAGCGGGCAGTTCGGTTTCAGGCAGGTCTTGCAACGTGACACCCTGTGCACGGCGGGAGATGCAATAGGTCAGGCTCTCGCTAAACTCCCCAATGTCAAGCACTTCCGGAATCGGGAGCGCGGCCGATGCAAAGTGCCGATAAACATAACGATCTTTGTAGAAACCATCGGCGCAGCTATTTACCCGCAGGACATATCCACGCCCTCCTACATCGAAGCTGAAAGCACGAGATTCTTCGCCCTCCGAGAGCTGCATCAGGTCGGAGACGCTGTCGAACTTTTCGATCAGAAACTTCTCGACAGACGTCGCGGTGAGTTCAGGCTTTTTCATATCTCATTGCCCCCCGGGATCTGCGAAAGCTCGAGAGAGATAGATTTGTAGAGAGAGACTGGTGATTTCAGCGTGTCCTCTCCAAATGAAATGAACTTCCTTATATAGAGGAAGGTCTTGCGAAGGATAGTGGGATTGTGCGTCATCCCTTACGTCAGTGGAGATATCACATCAATCCACTTGCTTTGAAGACGTGGTTGGAACGTCTTCTTTTTCCACGATGCTCCTCGTGGGTGGGGGTCCATCTTTGGGACCACTGTCGGCAGAGGCATCTTGAACGATAGCCTTTCCTTTATCGCAATGATGGCATTTGTAGGTGCCACCTTCCTTTTCTACTGTCCTTTTGATGAAGTGACAGATAGCTGGGCAATGGAATCCGAGGAGGTTTCCCGATATTACCCTTTGTTGAAAAGTCTCAATAGCCCTTTGGTCTTCTGAGACTGTATCTTTGATATTCTTGGAGTAGACGAGAGTGTCGTGCTCCACCATGTTATCACATCAATCCACTTGCTTTGAAGACGTGGTTGGAACGTCTTCTTTTTCCACGATGCTCCTCGTGGGTGGGGGTCCATCTTTGGGACCACTGTCGGCAGAGGCATCTTGAACGATAGCCTTTCCTTTATCGCAATGATGGCATTTGTAGGTGCCACCTTCCTTTTCTACTGTCCTTTTGATGAAGTGACAGATAGCTGGGCAATGGAATCCGAGGAGGTTTCCCGATATTACCCTTTGTTGAAAAGTCTCAATAGCCCTTTGGTCTTCTGAGACTGTATCTTTGATATTCTTGGAGTAGACGAGAGTGTCGTGCTCCACCATGTTGGCAAGCTGCTCTAGCCAATACGCAAACCGCCTCTCCCCGCGCGTTGGCCGATTCATTAATGCAGCTGGCACGACAGGTTTCCCGACTGGAAAGCGGGCAGTGAGCGCAACGCAATTAATGTGAGTTAGCTCACTCATTAGGCACCCCAGGCTTTACACTTTATGCTTCCGGCTCGTATGTTGTGTGGAATTGTGAGCGGATAACAATTTCACACAGGAAACAGCTATGACCATGATTACGCCAAGCTTTGAGACTTTTCAACAAAGGGTAATATCGGGAAACCTCCTCGGATTCCATTGCCCAGCTATCTGTCACTTCATCAAAAGGACAGTAGAAAAGGAAGGTGGCACCTACAAATGCCATCATTGCGATAAAGGAAAGGCTATCGTTCAAGATGCCTCTGCCGACAGTGGTCCCAAAGATGGACCCCCACCCACGAGGAGCATCGTGGAAAAAGAAGACGTTCCAACCACGTCTTCAAAGCAAGTGGATTGATGTGATAACATGGTGGAGCACGACACTCTCGTCTACTCCAAGAATATCAAAGATACAGTCTCAGAAGACCAAAGGGCTATTGAGACTTTTCAACAAAGGGTAATATCGGGAAACCTCCTCGGATTCCATTGCCCAGCTATCTGTCACTTCATCAAAAGGACAGTAGAAAAGGAAGGTGGCACCTACAAATGCCATCATTGCGATAAAGGAAAGGCTATCGTTCAAGATGCCTCTGCCGACAGTGGTCCCAAAGATGGACCCCCACCCACGAGGAGCATCGTGGAAAAAGAAGACGTTCCAACCACGTCTTCAAAGCAAGTGGATTGATGTGATATCTCCACTGACGTAAGGGATGACGCACAATCCCACTATCCTTCGCAAGACCTTCCTCTATATAAGGAAGTTCATTTCATTTGGAGAGGACACGCTGACTGCAGAAGCTTTAATACGACTCACTATAGGGAGACCACAACGGTTTCCCTAGAAATAATTTTGTTTAACTTTAAGAAGGAGATATAGGGCCCCCTAGGATGGCTACGACCGCCGCGGCCGCGGCCGCCGCCCTGTCCGCCGCCGCGACGGCCAAGACCGGCCGTAAGAACCACCAGCGACACCACGTCCTTCCCGCTCGAGGCCGGGTGGGGGCGGCGGCGGTCAGGTGCTCGGCGGTGTCCCCGGTCACCCCGCCGTCCCCGGCGCCGCCGGCCACGCCGCTCCGGCCGTGGGGGCCGGCCGAGCCCCGCAAGGGCGCGGACATCCTCGTGGAGGCGCTGGAGCGGTGCGGCGTCAGCGACGTGTTCGCCTACCCGGGCGGCGCGTCCATGGAGATCCACCAGGCGCTGACGCGCTCCCCGGTCATCACCAACCACCTCTTCCGCCACGAGCAGGGCGAGGCGTTCGCGGCGTCCGGGTACGCGCGCGCGTCCGGCCGCGTCGGGGTCTGCGTCGCCACCTCCGGCCCCGGGGCAACCAACCTCGTGTCCGCGCTCGCCGACGCGCTGCTCGACTCCGTCCCGATGGTCGCCATCACGGGCCAGGTCCCCCGCCGCATGATCGGCACCGACGCCTTCCAGGAGACGCCCATAGTCGAGGTCACCCGCTCCATCACCAAGCACAATTACCTTGTCCTTGATGTGGAGGACATCCCCCGCGTCATACAGGAAGCCTTCTTCCTCGCGTCCTCGGGCCGTCCTGGCCCGGTGCTGGTCGACATCCCCAAGGACATCCAGCAGCAGATGGCCGTGCCGGTCTGGGACACCTCGATGAATCTACCAGGGTACATCGCACGCCTGCCCAAGCCACCCGCGACAGAATTGCTTGAGCAGGTCTTGCGTCTGGTTGGCGAGTCACGGCGCCCGATTCTCTATGTCGGTGGTGGCTGCTCTGCATCTGGTGACGAATTGCGCTGGTTTGTTGAGCTGACTGGTATCCCAGTTACAACCACTCTGATGGGCCTCGGCAATTTCCCCAGTGACGACCCGTTGTCCCTGCGCATGCTTGGGATGCATGGCACGGTGTACGCAAATTATGCCGTGGATAAGGCTGACCTGTTGCTTGCGTTTGGTGTGCGGTTTGATGATCGTGTGACAGGGAAAATTGAGGCTTTTGCAAGCAGGGCCAAGATTGTGCACATTGACATTGATCCAGCAGAGATTGGAAAGAACAAGCAACCACATGTGTCAATTTGCGCAGATGTTAAGCTTGCTTTACAGGGCTTGAATGCTCTGCTACAACAGAGCACAACAAAGACAAGTTCTGATTTTAGTGCATGGCACAATGAGTTGGACCAGCAGAAGAGGGAGTTTCCTCTGGGGTACAAAACTTTTGGTGAAGAGATCCCACCGCAATATGCCATTCAGGTGCTGGATGAGCTGACGAAAGGTGAGGCAATCATCGCTACTGGTGTTGGGCAGCACCAGATGTGGGCGGCACAATATTACACCTACAAGCGGCCACGGCAGTGGCTGTCTTCGGCTGGTCTGGGCGCAATGGGATTTGGGCTGCCTGCTGCAGCTGGTGCTTCTGTGGCTAACCCAGGTGTCACAGTTGTTGATATTGATGGGGATGGTAGCTTCCTCATGAACATTCAGGAGCTGGCATTGATCCGCATTGAGAACCTCCCTGTGAAGGTGATGGTGTTGAACAACCAACATTTGGGTATGGTGGTGCAATGGGAGGATAGGTTTTACAAGGCGAATAGGGCGCATACATACTTGGGCAACCCGGAATGTGAGAGCGAGATATATCCAGATTTTGTGACTATTGCTAAGGGGTTCAATATTCCTGCAGTCCGTGTAACAAAGAAGAGTGAAGTCCGTGCCGCCATCAAGAAGATGCTCGAGACTCCAGGGCCATACTTGTTGGATATCATCGTCCCGCACCAGGAGCATGTGCTGCCTATGATCCCAAGTGGGGGCGCATTCAAGGACATGATCCTGGATGGTGATGGCAGGACTGTGTATTAAGGTACCTCTAGAACTAGCATAACCCCTTGGGGCCTCTAAACGGGTCTTGAGGGGTTTTTTGTTTCTCCATAATAATGTGTGAGTAGTTCCCAGATAAGGGAATTAGGGTTCCTATAGGGTTTCGCTCATGTGTTGAGCATATAAGAAACCCTTAGTATGTATTTGTATTTGTAAAATACTTCTATCAATAAAATTTCTAATTCCTAAAACCAAAATCCAGTACTAAAATCCAGATCTGCATGCCTGCAGGTCCACAAATTCGGGTCAAGGCGGAAGCCAGCGCGCCACCCCACGTCAGCAAATACGGAGGCGCGGGGTTGACGGCGTCACCCGGTCCTAACGGCGACCAACAAACCAGCCAGAAGAAATTACAGTAAAAAAAAAGTAAATTGCACTTTGATCCACCTTTTATTACCTAAGTCTCAATTTGGATCACCCTTAAACCTATCTTTTCAATTTGGGCCGGGTTGTGGTTTGGACTACCATGAACAACTTTTCGTCATGTCTAACTTCCCTTTCAGCAAACATATGAACCATATATAGAGGAGATCGGCCGTATACTAGAGCTGATGTGTTTAAGGTCGTTGATTGCACGAGAAAAAAAAATCCAAATCGCAACAATAGCAAATTTATCTGGTTCAAAGTGAAAAGATATGTTTAAAGGTAGTCCAAAGTAAAACTTATAGATAATAAAATGTGGTCCAAAGCGTAATTCACTCAAAAAAAATCAACGAGACGTGTACCAAACGGAGACAAACGGCATCTTCTCGAAATTTCCCAACCGCTCGCTCGCCCGCCTCGTCTTCCCGGAAACCGCGGTGGTTTCAGCGTGGCGGATTCTCCAAGCAGACGGAGACGTCACGGCACGGGACTCCTCCCACCACCCAACCGCCATAAATACCAGCCCCCTCATCTCCTCTCCTCGCATCAGCTCCACCCCCGAAAAATTTCTCCCCAATCTCGCGAGGCTCTCGTCGTCGAATCGAATCCTCTCGCGTCCTCAAGGTACGCTGCTTCTCCTCTCCTCGCTTCGTTTCGATTCGATTTCGGACGGGTGAGGTTGTTTTGTTGCTAGATCCGATTGGTGGTTAGGGTTGTCGATGTGATTATCGTGAGATGTTTAGGGGTTGTAGATCTGATGGTTGTGATTTGGGCACGGTTGGTTCGATAGGTGGAATCGTGGTTAGGTTTTGGGATTGGATGTTGGTTCTGATGATTGGGGGGAATTTTTACGGTTAGATGAATTGTTGGATGATTCGATTGGGGAAATCGGTGTAGATCTGTTGGGGAATTGTGGAACTAGTCATGCCTGAGTGATTGGTGCGATTTGTAGCGTGTTCCATCTTGTAGGCCTTGTTGCGAGCATGTTCAGATCTACTGTTCCGCTCTTGATTGAGTTATTGGTGCCATGGGTTGGTGCAAACACAGGCTTTAATATGTTATATCTGTTTTGTGTTTGATGTAGATCTGTAGGGTAGTTCTTCTTAGACATGGTTCAATTATGTAGCTTGTGCGTTTCGATTTGATTTCATATGTTCACAGATTAGATAATGATGAACTCTTTTAATTAATTGTCAATGGTAAATAGGAAGTCTTGTCGCTATATCTGTCATAATGATCTCATGTTACTATCTGCCAGTAATTTATGCTAAGAACTATATTAGAATATCATGTTACAATCTGTAGTAATATCATGTTACAATCTGTAGTTCATCTATATAATCTATTGTGGTAATTTCTTTTTACTATCTGTGTGAAGATTATTGCCACTAGTTCATTCTACTTATTTCTGAAGTTCAGGATACGTGTGCTGTTACTACCTATCTGAATACATGTGTGATGTGCCTGTTACTATCTTTTTGAATACATGTATGTTCTGTTGGAATATGTTTGCTGTTTGATCCGTTGTTGTGTCCTTAATCTTGTGCTAGTTCTTACCCTATCTGTTTGGTGATTATTTCTTGCAGATAGTTATCAGCAAGTTTGTACAAAGTGGT

**Sequences S6:** DNA sequence of the single-vector system **p35S-pT7-ALS/pUBI-AID-T7pol** (p35S_pT7_ALS_T7term_polA_pUBI_AID_T7pol_UGI_NOSterm)

TGATAACAGCGACTACAAGGATGACGATGACAAGGCTTAGAGCTCGAATTTCCCCGATCGTTCAAACATTTGGCAATAAAGTTTCTTAAGATTGAATCCTGTTGCCGGTCTTGCGATGATTATCATATAATTTCTGTTGAATTACGTTAAGCATGTAATAATTAACATGTAATGCATGACGTTATTTATGAGATGGGTTTTTATGATTAGAGTCCCGCAATTATACATTTAATACGCGATAGAAAACAAAATATAGCGCGCAAACTAGGATAAATTATCGCGCGCGGTGTCATCTATGTTACTAGATCGGGAATTCACTGGCCGTCGTTTTACACTGGCCGTCGTTTTACAACGTCGTGACTGGGAAAACCCTGGCGTTACCCAACTTAATCGCCTTGCAGCACATCCCCCTTTCGCCAGCTGGCGTAATAGCGAAGAGGCCCGCACCGATCGCCCTTCCCAACAGTTGCGCAGCCTGAATGGCGAATGCTAGAGCAGCTTGAGCTTGGATCAGATTGTCGTTTCCCGCCTTCAGTTTAAACTATCAGTGTTTGACAGGATATATTGGCGGGTAAACCTAAGAGAAAAGAGCGTTTATTAGAATAACGGATATTTAAAAGGGCGTGAAAAGGTTTATCCGTTCGTCCATTTGTATGTGCATGCCAACCACAGGGTTCCCCTCGGGATCAAAGTACTTTGATCCAACCCCTCCGCTGCTATAGTGCAGTCGGCTTCTGACGTTCAGTGCAGCCGTCTTCTGAAAACGACATGTCGCACAAGTCCTAAGTTACGCGACAGGCTGCCGCCCTGCCCTTTTCCTGGCGTTTTCTTGTCGCGTGTTTTAGTCGCATAAAGTAGAATACTTGCGACTAGAACCGGAGACATTACGCCATGAACAAGAGCGCCGCCGCTGGCCTGCTGGGCTATGCCCGCGTCAGCACCGACGACCAGGACTTGACCAACCAACGGGCCGAACTGCACGCGGCCGGCTGCACCAAGCTGTTTTCCGAGAAGATCACCGGCACCAGGCGCGACCGCCCGGAGCTGGCCAGGATGCTTGACCACCTACGCCCTGGCGACGTTGTGACAGTGACCAGGCTAGACCGCCTGGCCCGCAGCACCCGCGACCTACTGGACATTGCCGAGCGCATCCAGGAGGCCGGCGCGGGCCTGCGTAGCCTGGCAGAGCCGTGGGCCGACACCACCACGCCGGCCGGCCGCATGGTGTTGACCGTGTTCGCCGGCATTGCCGAGTTCGAGCGTTCCCTAATCATCGACCGCACCCGGAGCGGGCGCGAGGCCGCCAAGGCCCGAGGCGTGAAGTTTGGCCCCCGCCCTACCCTCACCCCGGCACAGATCGCGCACGCCCGCGAGCTGATCGACCAGGAAGGCCGCACCGTGAAAGAGGCGGCTGCACTGCTTGGCGTGCATCGCTCGACCCTGTACCGCGCACTTGAGCGCAGCGAGGAAGTGACGCCCACCGAGGCCAGGCGGCGCGGTGCCTTCCGTGAGGACGCATTGACCGAGGCCGACGCCCTGGCGGCCGCCGAGAATGAACGCCAAGAGGAACAAGCATGAAACCGCACCAGGACGGCCAGGACGAACCGTTTTTCATTACCGAAGAGATCGAGGCGGAGATGATCGCGGCCGGGTACGTGTTCGAGCCGCCCGCGCACGTCTCAACCGTGCGGCTGCATGAAATCCTGGCCGGTTTGTCTGATGCCAAGCTGGCGGCCTGGCCGGCCAGCTTGGCCGCTGAAGAAACCGAGCGCCGCCGTCTAAAAAGGTGATGTGTATTTGAGTAAAACAGCTTGCGTCATGCGGTCGCTGCGTATATGATGCGATGAGTAAATAAACAAATACGCAAGGGGAACGCATGAAGGTTATCGCTGTACTTAACCAGAAAGGCGGGTCAGGCAAGACGACCATCGCAACCCATCTAGCCCGCGCCCTGCAACTCGCCGGGGCCGATGTTCTGTTAGTCGATTCCGATCCCCAGGGCAGTGCCCGCGATTGGGCGGCCGTGCGGGAAGATCAACCGCTAACCGTTGTCGGCATCGACCGCCCGACGATTGACCGCGACGTGAAGGCCATCGGCCGGCGCGACTTCGTAGTGATCGACGGAGCGCCCCAGGCGGCGGACTTGGCTGTGTCCGCGATCAAGGCAGCCGACTTCGTGCTGATTCCGGTGCAGCCAAGCCCTTACGACATATGGGCCACCGCCGACCTGGTGGAGCTGGTTAAGCAGCGCATTGAGGTCACGGATGGAAGGCTACAAGCGGCCTTTGTCGTGTCGCGGGCGATCAAAGGCACGCGCATCGGCGGTGAGGTTGCCGAGGCGCTGGCCGGGTACGAGCTGCCCATTCTTGAGTCCCGTATCACGCAGCGCGTGAGCTACCCAGGCACTGCCGCCGCCGGCACAACCGTTCTTGAATCAGAACCCGAGGGCGACGCTGCCCGCGAGGTCCAGGCGCTGGCCGCTGAAATTAAATCAAAACTCATTTGAGTTAATGAGGTAAAGAGAAAATGAGCAAAAGCACAAACACGCTAAGTGCCGGCCGTCCGAGCGCACGCAGCAGCAAGGCTGCAACGTTGGCCAGCCTGGCAGACACGCCAGCCATGAAGCGGGTCAACTTTCAGTTGCCGGCGGAGGATCACACCAAGCTGAAGATGTACGCGGTACGCCAAGGCAAGACCATTACCGAGCTGCTATCTGAATACATCGCGCAGCTACCAGAGTAAATGAGCAAATGAATAAATGAGTAGATGAATTTTAGCGGCTAAAGGAGGCGGCATGGAAAATCAAGAACAACCAGGCACCGACGCCGTGGAATGCCCCATGTGTGGAGGAACGGGCGGTTGGCCAGGCGTAAGCGGCTGGGTTGTCTGCCGGCCCTGCAATGGCACTGGAACCCCCAAGCCCGAGGAATCGGCGTGACGGTCGCAAACCATCCGGCCCGGTACAAATCGGCGCGGCGCTGGGTGATGACCTGGTGGAGAAGTTGAAGGCCGCGCAGGCCGCCCAGCGGCAACGCATCGAGGCAGAAGCACGCCCCGGTGAATCGTGGCAAGCGGCCGCTGATCGAATCCGCAAAGAATCCCGGCAACCGCCGGCAGCCGGTGCGCCGTCGATTAGGAAGCCGCCCAAGGGCGACGAGCAACCAGATTTTTTCGTTCCGATGCTCTATGACGTGGGCACCCGCGATAGTCGCAGCATCATGGACGTGGCCGTTTTCCGTCTGTCGAAGCGTGACCGACGAGCTGGCGAGGTGATCCGCTACGAGCTTCCAGACGGGCACGTAGAGGTTTCCGCAGGGCCGGCCGGCATGGCCAGTGTGTGGGATTACGACCTGGTACTGATGGCGGTTTCCCATCTAACCGAATCCATGAACCGATACCGGGAAGGGAAGGGAGACAAGCCCGGCCGCGTGTTCCGTCCACACGTTGCGGACGTACTCAAGTTCTGCCGGCGAGCCGATGGCGGAAAGCAGAAAGACGACCTGGTAGAAACCTGCATTCGGTTAAACACCACGCACGTTGCCATGCAGCGTACGAAGAAGGCCAAGAACGGCCGCCTGGTGACGGTATCCGAGGGTGAAGCCTTGATTAGCCGCTACAAGATCGTAAAGAGCGAAACCGGGCGGCCGGAGTACATCGAGATCGAGCTAGCTGATTGGATGTACCGCGAGATCACAGAAGGCAAGAACCCGGACGTGCTGACGGTTCACCCCGATTACTTTTTGATCGATCCCGGCATCGGCCGTTTTCTCTACCGCCTGGCACGCCGCGCCGCAGGCAAGGCAGAAGCCAGATGGTTGTTCAAGACGATCTACGAACGCAGTGGCAGCGCCGGAGAGTTCAAGAAGTTCTGTTTCACCGTGCGCAAGCTGATCGGGTCAAATGACCTGCCGGAGTACGATTTGAAGGAGGAGGCGGGGCAGGCTGGCCCGATCCTAGTCATGCGCTACCGCAACCTGATCGAGGGCGAAGCATCCGCCGGTTCCTAATGTACGGAGCAGATGCTAGGGCAAATTGCCCTAGCAGGGGAAAAAGGTCGAAAAGCACTCTTTCCTGTGGATAGCACGTACATTGGGAACCCAAAGCCGTACATTGGGAACCGGAACCCGTACATTGGGAACCCAAAGCCGTACATTGGGAACCGGTCACACATGTAAGTGACTGATATAAAAGAGAAAAAAGGCGATTTTTCCGCCTAAAACTCTTTAAAACTTATTAAAACTCTTAAAACCCGCCTGGCCTGTGCATAACTGTCTGGCCAGCGCACAGCCGAAGAGCTGCAAAAAGCGCCTACCCTTCGGTCGCTGCGCTCCCTACGCCCCGCCGCTTCGCGTCGGCCTATCGCGGCCGCTGGCCGCTCAAAAATGGCTGGCCTACGGCCAGGCAATCTACCAGGGCGCGGACAAGCCGCGCCGTCGCCACTCGACCGCCGGCGCCCACATCAAGGCACCCTGCCTCGCGCGTTTCGGTGATGACGGTGAAAACCTCTGACACATGCAGCTCCCGGAGACGGTCACAGCTTGTCTGTAAGCGGATGCCGGGAGCAGACAAGCCCGTCAGGGCGCGTCAGCGGGTGTTGGCGGGTGTCGGGGCGCAGCCATGACCCAGTCACGTAGCGATAGCGGAGTGTATACTGGCTTAACTATGCGGCATCAGAGCAGATTGTACTGAGAGTGCACCATATGCGGTGTGAAATACCGCACAGATGCGTAAGGAGAAAATACCGCATCAGGCGCTCTTCCGCTTCCTCGCTCACTGACTCGCTGCGCTCGGTCGTTCGGCTGCGGCGAGCGGTATCAGCTCACTCAAAGGCGGTAATACGGTTATCCACAGAATCAGGGGATAACGCAGGAAAGAACATGTGAGCAAAAGGCCAGCAAAAGGCCAGGAACCGTAAAAAGGCCGCGTTGCTGGCGTTTTTCCATAGGCTCCGCCCCCCTGACGAGCATCACAAAAATCGACGCTCAAGTCAGAGGTGGCGAAACCCGACAGGACTATAAAGATACCAGGCGTTTCCCCCTGGAAGCTCCCTCGTGCGCTCTCCTGTTCCGACCCTGCCGCTTACCGGATACCTGTCCGCCTTTCTCCCTTCGGGAAGCGTGGCGCTTTCTCATAGCTCACGCTGTAGGTATCTCAGTTCGGTGTAGGTCGTTCGCTCCAAGCTGGGCTGTGTGCACGAACCCCCCGTTCAGCCCGACCGCTGCGCCTTATCCGGTAACTATCGTCTTGAGTCCAACCCGGTAAGACACGACTTATCGCCACTGGCAGCAGCCACTGGTAACAGGATTAGCAGAGCGAGGTATGTAGGCGGTGCTACAGAGTTCTTGAAGTGGTGGCCTAACTACGGCTACACTAGAAGGACAGTATTTGGTATCTGCGCTCTGCTGAAGCCAGTTACCTTCGGAAAAAGAGTTGGTAGCTCTTGATCCGGCAAACAAACCACCGCTGGTAGCGGTGGTTTTTTTGTTTGCAAGCAGCAGATTACGCGCAGAAAAAAAGGATCTCAAGAAGATCCTTTGATCTTTTCTACGGGGTCTGACGCTCAGTGGAACGAAAACTCACGTTAAGGGATTTTGGTCATGCATTCTAGGTACTAAAACAATTCATCCAGTAAAATATAATATTTTATTTTCTCCCAATCAGGCTTGATCCCCAGTAAGTCAAAAAATAGCTCGACATACTGTTCTTCCCCGATATCCTCCCTGATCGACCGGACGCAGAAGGCAATGTCATACCACTTGTCCGCCCTGCCGCTTCTCCCAAGATCAATAAAGCCACTTACTTTGCCATCTTTCACAAAGATGTTGCTGTCTCCCAGGTCGCCGTGGGAAAAGACAAGTTCCTCTTCGGGCTTTTCCGTCTTTAAAAAATCATACAGCTCGCGCGGATCTTTAAATGGAGTGTCTTCTTCCCAGTTTTCGCAATCCACATCGGCCAGATCGTTATTCAGTAAGTAATCCAATTCGGCTAAGCGGCTGTCTAAGCTATTCGTATAGGGACAATCCGATATGTCGATGGAGTGAAAGAGCCTGATGCACTCCGCATACAGCTCGATAATCTTTTCAGGGCTTTGTTCATCTTCATACTCTTCCGAGCAAAGGACGCCATCGGCCTCACTCATGAGCAGATTGCTCCAGCCATCATGCCGTTCAAAGTGCAGGACCTTTGGAACAGGCAGCTTTCCTTCCAGCCATAGCATCATGTCCTTTTCCCGTTCCACATCATAGGTGGTCCCTTTATACCGGCTGTCCGTCATTTTTAAATATAGGTTTTCATTTTCTCCCACCAGCTTATATACCTTAGCAGGAGACATTCCTTCCGTATCTTTTACGCAGCGGTATTTTTCGATCAGTTTTTTCAATTCCGGTGATATTCTCATTTTAGCCATTTATTATTTCCTTCCTCTTTTCTACAGTATTTAAAGATACCCCAAGAAGCTAATTATAACAAGACGAACTCCAATTCACTGTTCCTTGCATTCTAAAACCTTAAATACCAGAAAACAGCTTTTTCAAAGTTGTTTTCAAAGTTGGCGTATAACATAGTATCGACGGAGCCGATTTTGAAACCGCGGTGATCACAGGCAGCAACGCTCTGTCATCGTTACAATCAACATGCTACCCTCCGCGAGATCATCCGTGTTTCAAACCCGGCAGCTTAGTTGCCGTTCTTCCGAATAGCATCGGTAACATGAGCAAAGTCTGCCGCCTTACAACGGCTCTCCCGCTGACGCCGTCCCGGACTGATGGGCTGCCTGTATCGAGTGGTGATTTTGTGCCGAGCTGCCGGTCGGGGAGCTGTTGGCTGGCTGGTGGCAGGATATATTGTGGTGTAAACAAATTGACGCTTAGACAACTTAATAACACATTGCGGACGTTTTTAATGTACTGAATTAACGCCGAATTAATTCGGGGGATCTGGATTTTAGTACTGGATTTTGGTTTTAGGAATTAGAAATTTTATTGATAGAAGTATTTTACAAATACAAATACATACTAAGGGTTTCTTATATGCTCAACACATGAGCGAAACCCTATAGGAACCCTAATTCCCTTATCTGGGAACTACTCACACATTATTATGGAGAAACTCGAGCTTGTCGATCGACAGATCCGGTCGGCATCTACTCTATTTCTTTGCCCTCGGACGAGTGCTGGGGCGTCGGTTTCCACTATCGGCGAGTACTTCTACACAGCCATCGGTCCAGACGGCCGCGCTTCTGCGGGCGATTTGTGTACGCCCGACAGTCCCGGCTCCGGATCGGACGATTGCGTCGCATCGACCCTGCGCCCAAGCTGCATCATCGAAATTGCCGTCAACCAAGCTCTGATAGAGTTGGTCAAGACCAATGCGGAGCATATACGCCCGGAGTCGTGGCGATCCTGCAAGCTCCGGATGCCTCCGCTCGAAGTAGCGCGTCTGCTGCTCCATACAAGCCAACCACGGCCTCCAGAAGAAGATGTTGGCGACCTCGTATTGGGAATCCCCGAACATCGCCTCGCTCCAGTCAATGACCGCTGTTATGCGGCCATTGTCCGTCAGGACATTGTTGGAGCCGAAATCCGCGTGCACGAGGTGCCGGACTTCGGGGCAGTCCTCGGCCCAAAGCATCAGCTCATCGAGAGCCTGCGCGACGGACGCACTGACGGTGTCGTCCATCACAGTTTGCCAGTGATACACATGGGGATCAGCAATCGCGCATATGAAATCACGCCATGTAGTGTATTGACCGATTCCTTGCGGTCCGAATGGGCCGAACCCGCTCGTCTGGCTAAGATCGGCCGCAGCGATCGCATCCATAGCCTCCGCGACCGGTTGTAGAACAGCGGGCAGTTCGGTTTCAGGCAGGTCTTGCAACGTGACACCCTGTGCACGGCGGGAGATGCAATAGGTCAGGCTCTCGCTAAACTCCCCAATGTCAAGCACTTCCGGAATCGGGAGCGCGGCCGATGCAAAGTGCCGATAAACATAACGATCTTTGTAGAAACCATCGGCGCAGCTATTTACCCGCAGGACATATCCACGCCCTCCTACATCGAAGCTGAAAGCACGAGATTCTTCGCCCTCCGAGAGCTGCATCAGGTCGGAGACGCTGTCGAACTTTTCGATCAGAAACTTCTCGACAGACGTCGCGGTGAGTTCAGGCTTTTTCATATCTCATTGCCCCCCGGGATCTGCGAAAGCTCGAGAGAGATAGATTTGTAGAGAGAGACTGGTGATTTCAGCGTGTCCTCTCCAAATGAAATGAACTTCCTTATATAGAGGAAGGTCTTGCGAAGGATAGTGGGATTGTGCGTCATCCCTTACGTCAGTGGAGATATCACATCAATCCACTTGCTTTGAAGACGTGGTTGGAACGTCTTCTTTTTCCACGATGCTCCTCGTGGGTGGGGGTCCATCTTTGGGACCACTGTCGGCAGAGGCATCTTGAACGATAGCCTTTCCTTTATCGCAATGATGGCATTTGTAGGTGCCACCTTCCTTTTCTACTGTCCTTTTGATGAAGTGACAGATAGCTGGGCAATGGAATCCGAGGAGGTTTCCCGATATTACCCTTTGTTGAAAAGTCTCAATAGCCCTTTGGTCTTCTGAGACTGTATCTTTGATATTCTTGGAGTAGACGAGAGTGTCGTGCTCCACCATGTTATCACATCAATCCACTTGCTTTGAAGACGTGGTTGGAACGTCTTCTTTTTCCACGATGCTCCTCGTGGGTGGGGGTCCATCTTTGGGACCACTGTCGGCAGAGGCATCTTGAACGATAGCCTTTCCTTTATCGCAATGATGGCATTTGTAGGTGCCACCTTCCTTTTCTACTGTCCTTTTGATGAAGTGACAGATAGCTGGGCAATGGAATCCGAGGAGGTTTCCCGATATTACCCTTTGTTGAAAAGTCTCAATAGCCCTTTGGTCTTCTGAGACTGTATCTTTGATATTCTTGGAGTAGACGAGAGTGTCGTGCTCCACCATGTTGGCAAGCTGCTCTAGCCAATACGCAAACCGCCTCTCCCCGCGCGTTGGCCGATTCATTAATGCAGCTGGCACGACAGGTTTCCCGACTGGAAAGCGGGCAGTGAGCGCAACGCAATTAATGTGAGTTAGCTCACTCATTAGGCACCCCAGGCTTTACACTTTATGCTTCCGGCTCGTATGTTGTGTGGAATTGTGAGCGGATAACAATTTCACACAGGAAACAGCTATGACCATGATTACGCCAAGCTTTGAGACTTTTCAACAAAGGGTAATATCGGGAAACCTCCTCGGATTCCATTGCCCAGCTATCTGTCACTTCATCAAAAGGACAGTAGAAAAGGAAGGTGGCACCTACAAATGCCATCATTGCGATAAAGGAAAGGCTATCGTTCAAGATGCCTCTGCCGACAGTGGTCCCAAAGATGGACCCCCACCCACGAGGAGCATCGTGGAAAAAGAAGACGTTCCAACCACGTCTTCAAAGCAAGTGGATTGATGTGATAACATGGTGGAGCACGACACTCTCGTCTACTCCAAGAATATCAAAGATACAGTCTCAGAAGACCAAAGGGCTATTGAGACTTTTCAACAAAGGGTAATATCGGGAAACCTCCTCGGATTCCATTGCCCAGCTATCTGTCACTTCATCAAAAGGACAGTAGAAAAGGAAGGTGGCACCTACAAATGCCATCATTGCGATAAAGGAAAGGCTATCGTTCAAGATGCCTCTGCCGACAGTGGTCCCAAAGATGGACCCCCACCCACGAGGAGCATCGTGGAAAAAGAAGACGTTCCAACCACGTCTTCAAAGCAAGTGGATTGATGTGATATCTCCACTGACGTAAGGGATGACGCACAATCCCACTATCCTTCGCAAGACCTTCCTCTATATAAGGAAGTTCATTTCATTTGGAGAGGACACGCTGACTGCAGAAGCTTTAATACGACTCACTATAGGGAGACCACAACGGTTTCCCTAGAAATAATTTTGTTTAACTTTAAGAAGGAGATATAGGGCCCCCTAGGATGGCTACGACCGCCGCGGCCGCGGCCGCCGCCCTGTCCGCCGCCGCGACGGCCAAGACCGGCCGTAAGAACCACCAGCGACACCACGTCCTTCCCGCTCGAGGCCGGGTGGGGGCGGCGGCGGTCAGGTGCTCGGCGGTGTCCCCGGTCACCCCGCCGTCCCCGGCGCCGCCGGCCACGCCGCTCCGGCCGTGGGGGCCGGCCGAGCCCCGCAAGGGCGCGGACATCCTCGTGGAGGCGCTGGAGCGGTGCGGCGTCAGCGACGTGTTCGCCTACCCGGGCGGCGCGTCCATGGAGATCCACCAGGCGCTGACGCGCTCCCCGGTCATCACCAACCACCTCTTCCGCCACGAGCAGGGCGAGGCGTTCGCGGCGTCCGGGTACGCGCGCGCGTCCGGCCGCGTCGGGGTCTGCGTCGCCACCTCCGGCCCCGGGGCAACCAACCTCGTGTCCGCGCTCGCCGACGCGCTGCTCGACTCCGTCCCGATGGTCGCCATCACGGGCCAGGTCCCCCGCCGCATGATCGGCACCGACGCCTTCCAGGAGACGCCCATAGTCGAGGTCACCCGCTCCATCACCAAGCACAATTACCTTGTCCTTGATGTGGAGGACATCCCCCGCGTCATACAGGAAGCCTTCTTCCTCGCGTCCTCGGGCCGTCCTGGCCCGGTGCTGGTCGACATCCCCAAGGACATCCAGCAGCAGATGGCCGTGCCGGTCTGGGACACCTCGATGAATCTACCAGGGTACATCGCACGCCTGCCCAAGCCACCCGCGACAGAATTGCTTGAGCAGGTCTTGCGTCTGGTTGGCGAGTCACGGCGCCCGATTCTCTATGTCGGTGGTGGCTGCTCTGCATCTGGTGACGAATTGCGCTGGTTTGTTGAGCTGACTGGTATCCCAGTTACAACCACTCTGATGGGCCTCGGCAATTTCCCCAGTGACGACCCGTTGTCCCTGCGCATGCTTGGGATGCATGGCACGGTGTACGCAAATTATGCCGTGGATAAGGCTGACCTGTTGCTTGCGTTTGGTGTGCGGTTTGATGATCGTGTGACAGGGAAAATTGAGGCTTTTGCAAGCAGGGCCAAGATTGTGCACATTGACATTGATCCAGCAGAGATTGGAAAGAACAAGCAACCACATGTGTCAATTTGCGCAGATGTTAAGCTTGCTTTACAGGGCTTGAATGCTCTGCTACAACAGAGCACAACAAAGACAAGTTCTGATTTTAGTGCATGGCACAATGAGTTGGACCAGCAGAAGAGGGAGTTTCCTCTGGGGTACAAAACTTTTGGTGAAGAGATCCCACCGCAATATGCCATTCAGGTGCTGGATGAGCTGACGAAAGGTGAGGCAATCATCGCTACTGGTGTTGGGCAGCACCAGATGTGGGCGGCACAATATTACACCTACAAGCGGCCACGGCAGTGGCTGTCTTCGGCTGGTCTGGGCGCAATGGGATTTGGGCTGCCTGCTGCAGCTGGTGCTTCTGTGGCTAACCCAGGTGTCACAGTTGTTGATATTGATGGGGATGGTAGCTTCCTCATGAACATTCAGGAGCTGGCATTGATCCGCATTGAGAACCTCCCTGTGAAGGTGATGGTGTTGAACAACCAACATTTGGGTATGGTGGTGCAATGGGAGGATAGGTTTTACAAGGCGAATAGGGCGCATACATACTTGGGCAACCCGGAATGTGAGAGCGAGATATATCCAGATTTTGTGACTATTGCTAAGGGGTTCAATATTCCTGCAGTCCGTGTAACAAAGAAGAGTGAAGTCCGTGCCGCCATCAAGAAGATGCTCGAGACTCCAGGGCCATACTTGTTGGATATCATCGTCCCGCACCAGGAGCATGTGCTGCCTATGATCCCAAGTGGGGGCGCATTCAAGGACATGATCCTGGATGGTGATGGCAGGACTGTGTATTAAGGTACCTCTAGAACTAGCATAACCCCTTGGGGCCTCTAAACGGGTCTTGAGGGGTTTTTTGTTTCTCCATAATAATGTGTGAGTAGTTCCCAGATAAGGGAATTAGGGTTCCTATAGGGTTTCGCTCATGTGTTGAGCATATAAGAAACCCTTAGTATGTATTTGTATTTGTAAAATACTTCTATCAATAAAATTTCTAATTCCTAAAACCAAAATCCAGTACTAAAATCCAGATCTGCATGCCTGCAGGTCCACAAATTCGGGTCAAGGCGGAAGCCAGCGCGCCACCCCACGTCAGCAAATACGGAGGCGCGGGGTTGACGGCGTCACCCGGTCCTAACGGCGACCAACAAACCAGCCAGAAGAAATTACAGTAAAAAAAAAGTAAATTGCACTTTGATCCACCTTTTATTACCTAAGTCTCAATTTGGATCACCCTTAAACCTATCTTTTCAATTTGGGCCGGGTTGTGGTTTGGACTACCATGAACAACTTTTCGTCATGTCTAACTTCCCTTTCAGCAAACATATGAACCATATATAGAGGAGATCGGCCGTATACTAGAGCTGATGTGTTTAAGGTCGTTGATTGCACGAGAAAAAAAAATCCAAATCGCAACAATAGCAAATTTATCTGGTTCAAAGTGAAAAGATATGTTTAAAGGTAGTCCAAAGTAAAACTTATAGATAATAAAATGTGGTCCAAAGCGTAATTCACTCAAAAAAAATCAACGAGACGTGTACCAAACGGAGACAAACGGCATCTTCTCGAAATTTCCCAACCGCTCGCTCGCCCGCCTCGTCTTCCCGGAAACCGCGGTGGTTTCAGCGTGGCGGATTCTCCAAGCAGACGGAGACGTCACGGCACGGGACTCCTCCCACCACCCAACCGCCATAAATACCAGCCCCCTCATCTCCTCTCCTCGCATCAGCTCCACCCCCGAAAAATTTCTCCCCAATCTCGCGAGGCTCTCGTCGTCGAATCGAATCCTCTCGCGTCCTCAAGGTACGCTGCTTCTCCTCTCCTCGCTTCGTTTCGATTCGATTTCGGACGGGTGAGGTTGTTTTGTTGCTAGATCCGATTGGTGGTTAGGGTTGTCGATGTGATTATCGTGAGATGTTTAGGGGTTGTAGATCTGATGGTTGTGATTTGGGCACGGTTGGTTCGATAGGTGGAATCGTGGTTAGGTTTTGGGATTGGATGTTGGTTCTGATGATTGGGGGGAATTTTTACGGTTAGATGAATTGTTGGATGATTCGATTGGGGAAATCGGTGTAGATCTGTTGGGGAATTGTGGAACTAGTCATGCCTGAGTGATTGGTGCGATTTGTAGCGTGTTCCATCTTGTAGGCCTTGTTGCGAGCATGTTCAGATCTACTGTTCCGCTCTTGATTGAGTTATTGGTGCCATGGGTTGGTGCAAACACAGGCTTTAATATGTTATATCTGTTTTGTGTTTGATGTAGATCTGTAGGGTAGTTCTTCTTAGACATGGTTCAATTATGTAGCTTGTGCGTTTCGATTTGATTTCATATGTTCACAGATTAGATAATGATGAACTCTTTTAATTAATTGTCAATGGTAAATAGGAAGTCTTGTCGCTATATCTGTCATAATGATCTCATGTTACTATCTGCCAGTAATTTATGCTAAGAACTATATTAGAATATCATGTTACAATCTGTAGTAATATCATGTTACAATCTGTAGTTCATCTATATAATCTATTGTGGTAATTTCTTTTTACTATCTGTGTGAAGATTATTGCCACTAGTTCATTCTACTTATTTCTGAAGTTCAGGATACGTGTGCTGTTACTACCTATCTGAATACATGTGTGATGTGCCTGTTACTATCTTTTTGAATACATGTATGTTCTGTTGGAATATGTTTGCTGTTTGATCCGTTGTTGTGTCCTTAATCTTGTGCTAGTTCTTACCCTATCTGTTTGGTGATTATTTCTTGCAGATAGTTATCAGCAAGTTTGTACAAAAAAGCAGGCTTCGAAGGAGATAGAACCAATTCTCTAAGGAAATACTTAACCATGGACTATAAGGACCACGACGGAGACTACAAGGATCATGATATTGATTACAAAGACGATGACGATAAGATGGCCCCAAAGAAGAAGCGGAAGGTGGGTATCCACGGAGTCCCAGCAGCCGCATGCCCCGGGATGGACAGCCTCTTGATGAACCGGAGGGAGTTTCTTTACCAATTCAAAAATGTCCGCTGGGCTAAGGGTCGGCGTGAGACCTACCTGTGCTACGTAGTGAAGAGGCGTGACAGTGCTACATCCTTTTCACTGGACTTTGGTTATCTTCGCAATAAGAACGGCTGCCACGTGGAATTGCTCTTCCTCCGCTACATCTCGGACTGGGACCTAGACCCTGGCCGCTGCTACCGCGTCACCTGGTTCATCTCCTGGAGCCCCTGCTACGACTGTGCCCGACATGTGGCCGACTTTCTGCGAGGGAACCCCAACCTCAGTCTGAGGATCTTCACCGCGCGCCTCTACTTCTGTGAGGACCGCAAGGCTGAGCCCGAGGGGCTGCGGCGGCTGCACCGCGCCGGGGTGCAAATAGCCATCATGACCTTCAAAGATTATTTTTACTGCTGGAATACTTTTGTAGAAAACCACGGAAGAACTTTCAAAGCCTGGGAAGGGCTGCATGAAAATTCAGTTCGTCTCTCCAGACAGCTTCGGCGCATCCTTTTGCCCCTGTATGAGGTTGATGACTTACGAGACGCATTTCGTACTAGCGGCAGCGAGACTCCCGGGACCTCAGAGTCCGCCACACCCGAAAGTAACACCATCAACATTGCTAAGAACGACTTCTCAGACATAGAGCTCGCGGCTATTCCGTTCAACACCCTGGCTGACCACTACGGCGAGAGACTCGCTAGGGAGCAGCTGGCGTTGGAGCATGAATCCTACGAGATGGGCGAGGCTAGGTTCCGCAAGATGTTCGAGCGACAATTGAAGGCAGGGGAGGTGGCGGACAACGCTGCCGCCAAGCCCCTGATCACAACCTTGCTGCCCAAAATGATCGCGCGGATCAACGATTGGTTTGAGGAGGTTAAGGCAAAACGGGGCAAACGCCCGACCGCATTTCAATTCCTCCAAGAAATCAAGCCTGAGGCTGTTGCCTACATCACTATCAAGACGACACTGGCGTGTCTCACAAGCGCCGACAACACCACCGTGCAAGCCGTCGCCAGCGCCATCGGGCGGGCAATTGAGGATGAGGCACGGTTTGGTAGGATCCGAGACCTGGAAGCGAAGCACTTCAAGAAGAACGTGGAAGAGCAGTTGAACAAACGCGTCGGCCACGTGTATAAAAAGGCTTTCATGCAGGTGGTGGAGGCCGATATGCTCAGTAAGGGGCTGCTTGGGGGGGAGGCGTGGTCATCCTGGCACAAGGAGGATAGCATTCACGTGGGGGTCCGATGTATCGAGATGCTGATAGAGAGCACCGGAATGGTCTCCCTCCATCGCCAGAACGCTGGGGTCGTAGGGCAGGACTCCGAGACTATTGAGCTGGCCCCCGAGTATGCCGAAGCAATCGCTACACGCGCAGGTGCACTGGCTGGGATAAGCCCTATGTTTCAGCCCTGCGTAGTGCCTCCAAAGCCATGGACCGGCATCACAGGGGGTGGCTATTGGGCCAACGGTAGGCGGCCTCTGGCCCTGGTACGCACGCACAGCAAGAAGGCGCTCATGCGCTATGAAGACGTTTACATGCCCGAGGTTTACAAGGCGATCAATATCGCGCAGAACACCGCCTGGAAAATCAATAAGAAGGTGTTGGCGGTCGCAAACGTGATTACCAAGTGGAAGCATTGCCCAGTCGAGGACATACCCGCCATAGAACGCGAAGAGCTGCCGATGAAGCCGGAAGACATTGATATGAACCCCGAGGCCCTCACCGCGTGGAAAAGAGCCGCAGCCGCCGTATACAGGAAGGATAAAGCGCGCAAGTCCCTACGCATAAGCCTCGAGTTTATGCTGGAACAGGCCAACAAGTTCGCCAACCACAAAGCTATCTGGTTCCCCTACAACATGGACTGGAGAGGGAGGGTCTACGCCGTCAGCATGTTCAATCCCCAGGGCAACGACATGACGAAGGGCCTTCTGACATTGGCAAAGGGGAAGCCTATCGGAAAGGAGGGGTACTACTGGCTCAAGATCCACGGCGCCAACTGCGCGGGAGTGGACAAGGTTCCATTTCCCGAGCGAATTAAGTTCATCGAGGAAAACCACGAAAACATTATGGCGTGCGCTAAATCCCCCCTCGAGAACACATGGTGGGCCGAGCAAGACTCCCCGTTCTGTTTTTTGGCATTCTGCTTTGAGTACGCCGGTGTGCAGCACCATGGCCTCTCATACAACTGTTCCCTGCCCCTGGCCTTCGACGGAAGTTGCAGTGGGATTCAACATTTCAGCGCAATGTTGCGGGACGAGGTCGGTGGCAGGGCCGTTAACCTGCTCCCTTCCGAAACGGTGCAGGACATCTACGGAATCGTGGCAAAAAAGGTAAACGAGATCCTGCAAGCGGATGCCATCAACGGGACGGACAATGAGGTCGTTACGGTGACAGACGAAAATACTGGGGAAATAAGCGAAAAGGTCAAGCTGGGGACCAAAGCACTCGCGGGTCAGTGGCTCGCCTACGGGGTGACACGCTCCGTCACCAAGAGAAGCGTGATGACCCTCGCGTACGGTTCAAAAGAATTCGCCTTCCGCCAGCAAGTGCTGGAGGACACCATCCAGCCGGCGATTGACTCCGGGAAGGGTCTCATGTTTACCCAGCCGAACCAGGCCGCAGGGTACATGGCCAAACTGATCTGGGAAAGCGTTAGCGTCACAGTGGTCGCCGCGGTTGAGGCGATGAATTGGCTGAAGAGCGCGGCAAAGCTCCTCGCCGCTGAGGTGAAGGACAAAAAGACCGGCGAAATCCTGCGCAAGCGCTGCGCCGTCCACTGGGTCACGCCGGATGGATTCCCCGTCTGGCAGGAGTACAAGAAGCCCATCCGAACCCGGCTCAACTTGATGTTCCTTGGCCAGTTTCGCCTGCAGCCCACGATAAACACCAACAAAGACAGCGAGATCGACGCCCACAAGCAGGAGAGCGGCATCGCGCCCAACTTCGTGCACAGTCAGGACGGGTCCCATCTGCGGAAAACTGTTGTGTGGGCTCACGAGAAGTACGGCATTGAGAGCTTCGCCCTGATACACGACAGCTTCGGGACCATACCAGCGGACGCAGCGAACCTGTTCAAAGCCGTGCGGGAAACAATGGTCGACACCTACGAAAGCTGCGACGTACTGGCAGACTTCTATGACCAATTCGCCGACCAGCTTCACGAGTCACAGCTCGACAAGATGCCCGCTCTGCCCGCGAAAGGCAACCTGAATTTGCGCGACATCCTTGAGAGCGATTTTGCGTTCGCCTCTGGTGGTTCTACTAATCTGTCAGATATTATTGAAAAGGAGACCGGTAAGCAACTGGTTATCCAGGAATCCATCCTCATGCTCCCAGAGGAGGTGGAAGAAGTCATTGGGAACAAGCCGGAAAGCGATATACTCGTGCACACCGCCTACGACGAGAGCACCGACGAGAATGTCATGCTTCTGACTAGCGACGCCCCTGAATACAAGCCTTGGGCTCTGGTCATACAGGATAGCAACGGTGAGAACAAGATTAAGATGCTCTCTGGTGGTTCTCCCAAGAAGAAGAGGAAAGTCATGCATTAGACCCAGCTTTCTTGTACAAAGTGGT
